# An original model of brain infection identifies the hijacking of host lipoprotein import as a bacterial strategy for blood-brain barrier crossing

**DOI:** 10.1101/2020.02.28.970376

**Authors:** Billel Benmimoun, Florentia Papastefanaki, Bruno Périchon, Katerina Segklia, Nicolas Roby, Vivi Miriagou, Christine Schmitt, Shaynoor Dramsi, Rebecca Matsas, Pauline Spéder

## Abstract

Pathogens able to cross the blood-brain barrier (BBB) induce long-term neurological sequelae and death. Understanding how neurotropic pathogens bypass this strong physiological barrier is a prerequisite to devise therapeutic strategies. Here we propose an innovative model of infection in the developing Drosophila brain, combining whole brain explants with *in vivo* systemic infection. We identified several mammalian pathogens able to cross the Drosophila BBB, including Group B Streptococcus (GBS). Amongst GBS surface components, lipoproteins, and in particular the B leucin-rich Blr, were important for BBB crossing and virulence in Drosophila. Further, we identified (V)LDL receptor LpR2, expressed in the BBB, as a host receptor for Blr, allowing GBS translocation through endocytosis. Finally, we demonstrated that Blr is required for BBB crossing and pathogenicity in a murine model of infection. Our results support the relevance of Drosophila for studying host-pathogen interactions and identify a new mechanism by which pathogens exploit host barriers to generate brain infection.

## Introduction

Central nervous system (CNS) infections are rare, yet extremely damaging. They lead to fatal outcomes and long-term neurological disabilities in surviving infants and adults, including cognitive deficit and motor impairment^1^. CNS infections are caused by the entry of pathogenic agents – bacteria, fungi or viruses, from the systemic environment into the CNS, provoking neuroinflammation and cellular damages.

A major route for CNS infection is the bloodstream, which pathogens enter after crossing the epithelial barriers of the skin and gut^2–4^, and in which they circulate as free particles or carried by blood cells^5^. To infect the brain, pathogens must ultimately bypass an additional guardian: the blood-brain barrier (BBB)^6^. The BBB is both a selective physical and chemical filter, enabling neuroprotective functions^7^. In higher vertebrates, brain microvascular endothelial cells form the core structure of the BBB. These cells are equipped to provide selective insulation, harboring intercellular tight junctions, absence of fenestrae, and asymmetrically localised transport systems^8,9^. These unique features distinguish brain microvascular endothelial cells from peripheral endothelial cells, and allow them to control ion, nutrient, and hormone transport ensuring ionic homeostasis and neuronal functions^10,11^. The BBB also includes perivascular pericytes, astrocytes and a basal membrane made of extracellular matrix, which regulate BBB integrity and endothelial functions^8,12^. This complex set of interlinked layers behaves as a double-edge sword for the organism: it restricts the entry of pathogens as well as therapeutic molecules, such as antibiotics^13,14^.

Pathogens that manage to cross the BBB thus secure their access to the CNS, where they tend to be immunologically-protected. Accordingly, neuro-invasive, neurotropic pathogens have developed intricate mechanisms that allow them to cross this layer and invade the CNS^2,3,6^. Three main strategies have been proposed so far: transcellular, paracellular and Trojan horse. The transcellular entry occurs through a receptor-mediated mechanism or pinocytosis, while the paracellular mechanism follows the increase of BBB permeability due to tight junction disruption. The Trojan horse mechanism uses infected blood cells which transmigrate from the periphery to the CNS. It has been proposed that pathogens could actually use several of these routes to invade the brain^6^.

So far, most of this knowledge comes from *in vitro* models of BBB^15,16^ where a monolayer of endothelial cells are co-cultured with pericytes and astrocytes in transwells^17^. However, these models still display a lot of variations in their tightness depending on the endothelial cells used and thus reproducibility is a major issue. Despite induced pluripotent stem cell-related advances ^17^ and new set-ups like microfluidic organ-on-chips^18^, these models struggle to recapitulate complex parameters crucial to BBB properties, including 3D architecture and dynamic cellular interactions. Animal models, mostly mice and rats, but also zebrafish, exist and have provided essential contributions to mechanistic explorations^3,16^, including revisiting results from *in vitro* models^19^. Manipulating these organisms to reach cellular resolution and causal relationships is nevertheless still highly challenging. Cost and ethical issues also hinder their extensive use.

Drosophila is a powerful and tractable model system, with unrivalled genetics. It has been very successful in identifying conserved molecular mechanisms in innate immunity, such as the Toll pathway^20–22^. Most studies focused on systemic and epithelial immunity (skin and gut)^23,24^, albeit one elegant study assessed the CNS inflammation mechanism during Zika virus infection in adults^25^. Strikingly, many aspects of mammalian neurogenesis are conserved in the Drosophila larva CNS, a post-embryonic, juvenile stage which also harbours a BBB (Fig. 1a-b). The open circulatory system of the fly, powered by the cardiac tube, carries the hemolymph, which is in direct contact with all the organs including the CNS. The BBB represents its outermost structure. It is exclusively of glial nature, a property found in invertebrates but also in lower vertebrates, such as sharks^26^, and is composed of two cell layers^27,28^. The subperineurial glia (SPG) are large polarised cells forming an epithelium-like structure with septate junctions (Fig. 1c), the equivalent of tight junctions in vertebrates. These represent a physical barrier to paracellular diffusion, similarly to the mammalian brain vascular endothelium^11,29^. The perineurial glia (PG), which do not have septate junctions, cover the SPG and are proposed to be a hemolymph sensor^30,31^. Transcriptional analysis of the SPG and PG layers uncovered a striking conservation of molecules with mouse BBB cells, including transporters and cell adhesion molecules ^32^. A recent study has confirmed the relevance of the fly model, showing shared mechanisms of response to xenobiotics^33^. Thus, the Drosophila BBB represents a physical and chemical barrier that retains conserved chemoprotective strategies with the mammalian BBB, ensuring brain homeostasis and protecting the brain from toxins and pathogens^34^.

**Figure 1:**
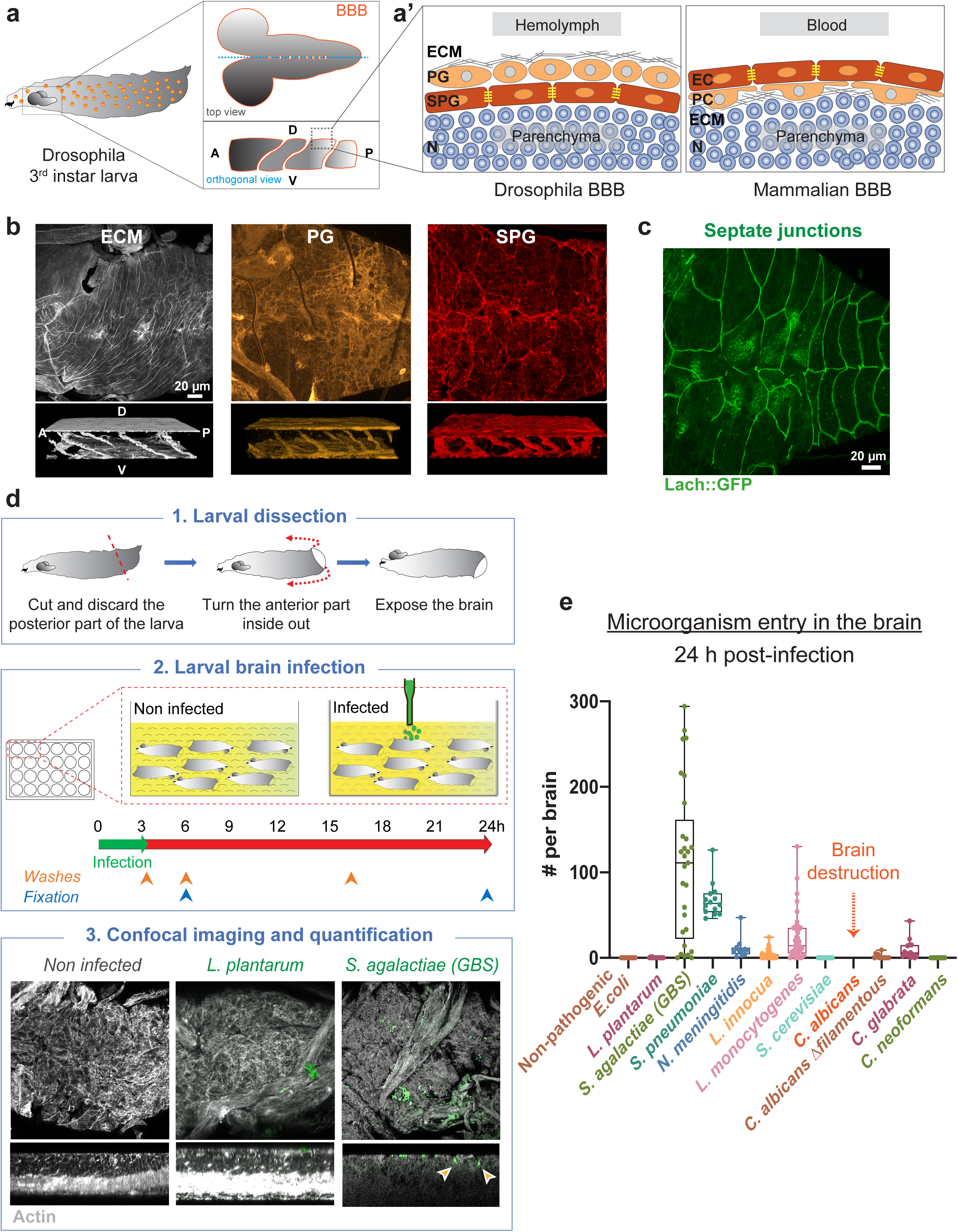
Screening for mammalian neuro-invasive pathogens in a brain explant set-up identifies Group B Streptococcus as able to cross the Drosophila blood-brain barrier. a. Schematic representation of Drosophila third instar larva showing the brain (grey) suspended in the hemolymph (orange). Top and orthogonal views of the brain covered by the blood-brain barrier (BBB) in dark orange. A’. Schematic representations of the composite Drosophila and mammalian BBBs, which include a layer of extracellular matrix (ECM in grey), a regulatory layer (perineurial glia (PG) and pericytes (PC) in light orange) and a barrier layer (subperineurial glia (SPG) and endothelial cells (EC) in dark orange) harbouring strong cell junctions (septate junctions and tight junctions in yellow). Brain cell populations, including neurons (N) are illustrated in blue. A, anterior. P, posterior. D, dorsal. V, ventral. b. Confocal images of the Drosophila BBB (top view and 3D orthogonal view) labelled for the ECM in grey (*vkg::GFP*), the PG in light orange (*NP6293-GAL4>mCD8-GFP*) and the SPG in red (*mdr65-mtd-tomato*). c. Septate junctions in green (*Lachesin::GFP*). d. Sketch of the *ex vivo* protocol to generate infection in Drosophila larval brains. Step 3 depicts confocal images of the brains (top and orthogonal views) stained with phalloidin (white). Pathogens (*L. plantarum* and *S. agalactiae [GBS]*) are stained in green. Orange arrows show GBS inside the brain parenchyma. e. Screening mammalian neurotropic pathogens for their ability to cross the Drosophila BBB and invade the brain. *S. agalactiae (GBS)* (n = 29), *S. pneumoniae* (n = 14), *N. meningitidis* (n = 11), *L. monocytogenes* (n = 43), and *C. glabrata* (n = 14) were able to cross the Drosophila BBB. *C. albicans* exists in a filamentous, hyphal form linked to pathogenicity, which destroyed the BBB and brain cells (n = 12), and in a yeast, non-hyphal form (*C. albicans Δfilamentous*) which was able to enter the brain (n = 14). *L. innocua* (n = 53) rarely crossed the BBB, while non-pathogenic *E. coli* (n = 12), *L. plantarum* (n = 14), *S. cerevisiae* and *C. neoformans* (n = 11) were not able to invade the brain.

Here we show that the Drosophila larval brain is a relevant and valuable system to model brain infection and discover cellular mechanisms of BBB crossing by mammalian pathogens. Taking advantage of this innovative model, we identified Blr, a lipoprotein displaying leucine rich repeat domains, as a new virulence factor contributing to GBS neurotropism in the fly and mouse. We further identified the Drosophila lipoprotein receptor LpR2 as a host receptor for Blr in the BBB, mediating GBS internalisation through endocytosis.

## Results

### Group B Streptococcus actively invades the Drosophila larval brain in an explant set up

Establishing a model of brain infection in the fly larva required developing experimental set-ups in which whole, intact living brains would be in contact with selected pathogens. We first devised an *ex vivo* protocol, as a straightforward, versatile platform for screening pathogens and conditions (Fig. 1d). Whole third instar larvae were opened posteriorly to expose all tissues while preventing damages to the brain and minimising injuries of the peripheral nerves. These brain explants, (including all larval tissues minus the gut, see Methods) were transferred to culture conditions that preserve cell viability, cell proliferation and BBB permeability, and that does not induce oxidative stress (Supp. Fig. 1a-d). In these conditions, brain explants could be kept for up to 48 h, at 30°C. The culture medium was then inoculated with selected pathogens at selected doses (see Methods). Brain explants were left in contact with pathogens for a given time (usually 3 h) to allow binding, washed several times to remove unattached microorganisms and kept in culture until the desired time of analysis. Whole fixed brains were scanned via confocal microscopy, allowing precise localisation and quantification of individual pathogens to assess brain entry and distinguish it from adhesion (Fig. 1d).

We used this set-up to screen for prokaryotic or eukaryotic pathogens known to trigger encephalitis and/or meningitis in mammals. We found that several were able to cross the Drosophila BBB and generate brain infection 24 h after inoculation (*Streptococcus agalactiae, Streptococcus pneumoniae, Neisseria meningitidis, Listeria monocytogenes, Candida glabrata* and non-hyphal *Candida albicans*; Fig. 1e). In contrast, non-pathogenic strains (*Lactobacillus plantarum*, non-pathogenic *Escherichia coli*, and *Saccharomyces cerevisiae*) were not able to enter the Drosophila brain, pointing to specific entry mechanisms under these conditions.

Amongst the various pathogens tested, *S. agalactiae* (Group B Streptococcus, GBS) proved to be the most efficient to cross the fly BBB and was detected both inside the brain and attached to its surface (Fig. 2a). We thus focused on GBS.

**Figure 2:**
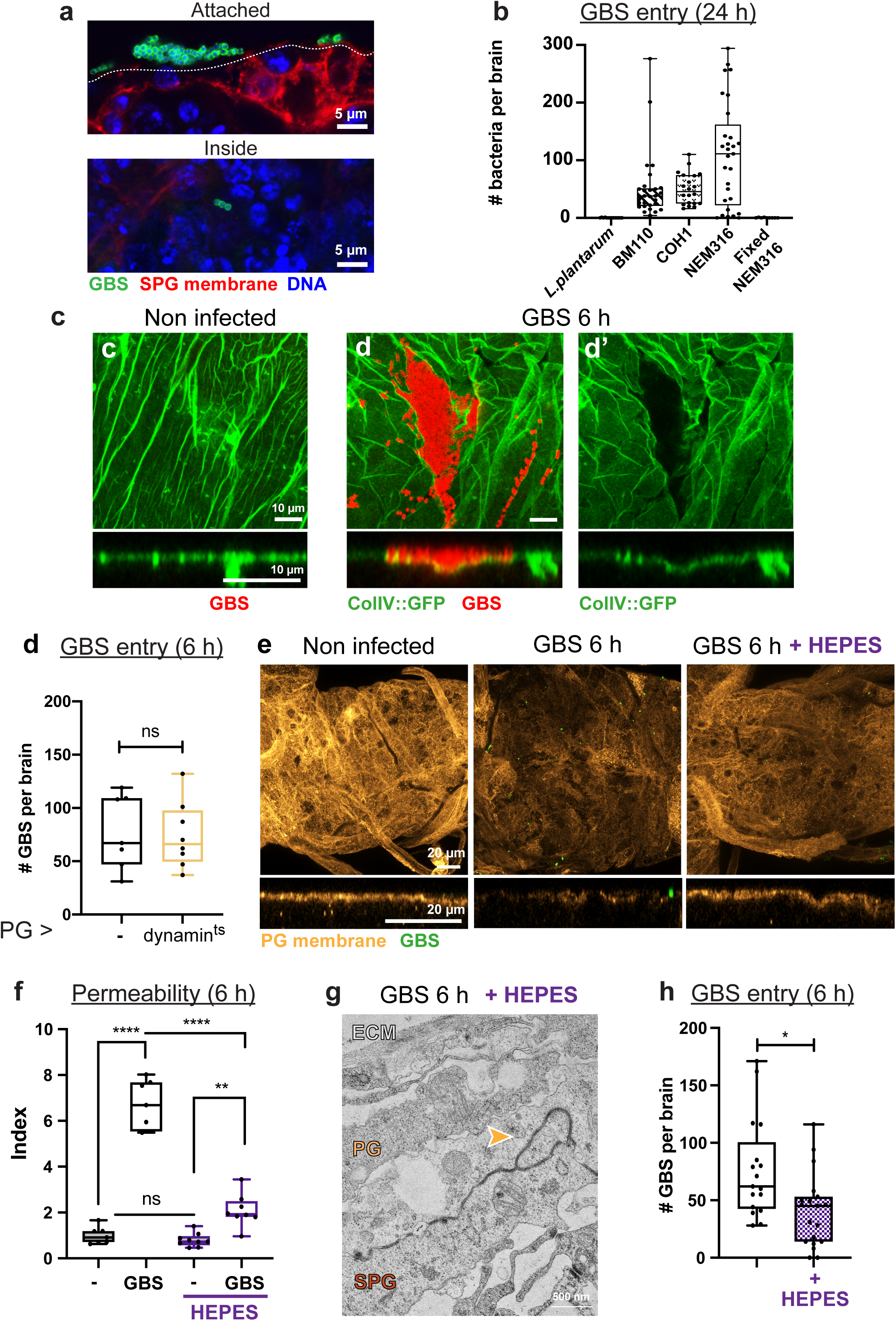
GBS uses a panel of strategies to cross the multiple layers of the BBB. a. Close-up of GBS (anti-GBS, green) attached to the SPG (*mdr65-mtd-Tomato*, red) as well as inside the brain. DAPI is in blue. b. Bacterial count inside the brain 24 h post infection for *L. plantarum* (n = 14), *GBS BM110* strain (n = 29), *GBS COH1* strain (n = 20), *GBS NEM316* strain (n = 29), and fixed *GBS NEM316* strain (n = 9). c. Confocal images (top view and close up orthogonal view) of non-infected and GBS-infected brains at 6 h post infection showing Collagen IV staining (*vkg::GFP*, green) and GBS (red). d. GBS entry at 6 h post infection was not significantly changed when endocytosis was blocked (dynamin^ts^) specifically in the PG. t Student’s t test: p(control vs *PG>dynamin*^*ts*^) = 0.8484. Control (n = 7); *PG>dynamin*^*ts*^ (n = 8). e. Confocal images (top view and close up orthogonal view) of non-infected and of GBS-infected brains without and with HEPES at 6 h post-GBS infection, showing PG membrane (*NP6293-GAL4>mCD8-GFP*, light orange). GBS, green. f. BBB permeability tests for non-infected (-) and GBS-infected brains at 6 h post infection without (black) and with HEPES (purple). ANOVA test: p(-vs GBS WT) < 0.0001; p(-HEPES vs GBS WT HEPES) = 0.0022; p (GBS vs GBS+HEPES) < 0.0001. n(-) = 7, n(GBS WT) = 7, n(-HEPES) = 8, n(GBS WT+HEPES) = 8. g. Transmission electron microscopy (TEM) picture of a Drosophila brain infected by GBS 6 h post infection, showing the different BBB layers and septate junctions (orange arrow). h. Bacterial count inside the brain in GBS entry without (black) and with HEPES (purple) at 6 h (Mann-Whitney test, p = 0.0102). n(GBS) = 17 and n(GBS+HEPES) = 19.

GBS is an opportunistic gram-positive bacterium responsible for severe invasive infections in neonates developing in pneumonia, septicaemia and meningitis^35–37^. Despite available antibiotic treatments and intrapartum prophylaxis, these cases still represent 10% of mortality and neurological sequelae in 25 to 50% of survivors, including cognitive impairment, seizures, hearing loss, and blindness^38,39^. Among the various GBS clinical isolates tested, NEM316 was the most efficient to infect Drosophila larval brain explants, and chosen as our reference GBS strain (Fig. 2b, from now on called GBS). Interestingly, dead, formaldehyde-fixed GBS were unable to enter Drosophila brain explants, underlining the fact that GBS needs to be alive and active to cross the BBB.

### Dissecting the panel of strategies used by GBS to cross the multiple layers of the BBB

The BBB is a composite structure with several protective layers. In Drosophila, GBS has first to overcome a layer of extracellular matrix (ECM) and then the PG (Fig. 1a-b). Indeed, we found GBS embedded in a protein trap (*vkg::GFP*^40^) for collagen IV, a major, conserved ECM component of the extracellular matrix (Fig. 2c). Whereas a collagen IV layer was still present under GBS infection, it appeared disturbed, generally weaker and sometimes absent where the bacteria were detected. Interestingly, GBS displays lytic activity on a collageno-mimetic peptide (Jackson et al., 1994), suggesting that GBS could destroy the ECM to gain access to the cellular layers. The first cellular layer, the PG, does not have intercellular junctions and thus a paracellular route could be used. To determine whether GBS relies on internalisation to cross this layer, we blocked endocytosis specifically in the PG by preventing dynamin (Drosophila Shibire) function through the overexpression of a temperature-sensitive (*shibire*^*ts*^) non-functional form. Interestingly, we found that preventing endocytosis in this layer did not alter brain invasion (Fig. 2d). Moreover, staining for a PG membrane reporter revealed a partial and much fainter signal under infection (Fig. 2e). These alterations in PG structure were not particularly associated with GBS localisation, but rather brain-wide, suggesting a systemic origin. Interestingly, extracellular acidosis has been shown to build in the brain under meningitis^41,42^, and GBS is known to secrete lactic acid and acidify the culture environment^39^. Measuring the pH of the culture medium at 3 h post infection actually revealed a strong acidification (Supp. Fig. 2c). To assess the impact of acidification to PG disruption, we buffered the cultured medium with HEPES. Blocking medium acidification nearly completely restored PG structure (Fig. 2e). Taking these results together, we propose that disruption of the ECM precedes the bacterial traversal of the PG layer, through paracellular mechanisms or acidosis-dependent destruction.

Ultimately, the capacity of GBS to invade the brain and generate infection is linked to its ability to cross the physical component of the BBB, harbouring cellular junctions^3,37^. This corresponds to the SPG in Drosophila. We first assayed BBB permeability of brain explants by dextran diffusion (see Methods), and found that it was significantly increased during GBS infection (Fig. 2f). We also noticed, in some cases, an altered morphology of SPG membrane and septate junctions, as assessed by the use of specific markers at confocal microscopy resolution (Supp. Fig. 2a-b). We assessed the contribution of acidification to these changes in BBB parameters, and found that blocking medium acidification strongly prevented the increase in BBB permeability observed under GBS infection (Fig. 2f), and restored BBB and septate junction morphologies (Fig. 2g and Supp. Fig. 2a-b and 2d-e). Bacterial counts in the brain also significantly changed in buffered medium compared to non-buffered (Fig. 2h). However, this change was moderate, and furthermore we were able to detect bacteria attached to the brain surface through scanning electron microscopy (SEM, Supp. Fig. 2f). Altogether these data show that, although acidification might indeed help GBS by altering BBB features, it is not a prerequisite for brain entry. This argues for the critical involvement of specific mechanisms for SPG crossing by GBS, on top of acidity-induced host tissue alteration.

### The B Streptoccocal surface lipoprotein Blr is required for BBB crossing

To identify GBS surface component(s) involved in this process, we first tested known factors implicated in pathogen virulence and adhesion to endothelial cells (Fig. 3a-a’), such as the polysaccharide capsule (acapsular mutant Δ*cpsE*), the hemolytic lipid toxin (non-hemolytic strain Δ*cylE* and hyper-hemolytic strain *cyl+*), or cell-wall anchored proteins (Δ*srtA*). None of these mutants were strongly affecting BBB crossing (Suppl. Fig. 3a). We thus tested the contribution of lipoproteins which are tethered to the cell membrane by an N-terminal lipid moiety. In Gram^+^ bacteria, lipoprotein biosynthesis involves two specific enzymes, i) Lgt (prolipoproteindiacylglyceryltransferase),whichattaches thelipidanchortothe prolipoproteins and ii) Lsp (lipoprotein signal peptidase), which specifically removes the N-terminal signal peptide of fully mature lipoproteins. In our model, removing either Lgt or Lsp decreased bacterial count within the brain at 24 h post infection compared to wild-type (WT) GBS, and the double mutant (Δ*lgt/lsp*) displayed an additive drop (Fig. 3b), leading to a strong impairment in GBS brain entry. GBS translocation into the brain was next assessed at earlier time points and a significant decrease was also demonstrated at 6 h post-infection for *Δlgt/lsp* mutant (Fig. 3c).

**Figure 3:**
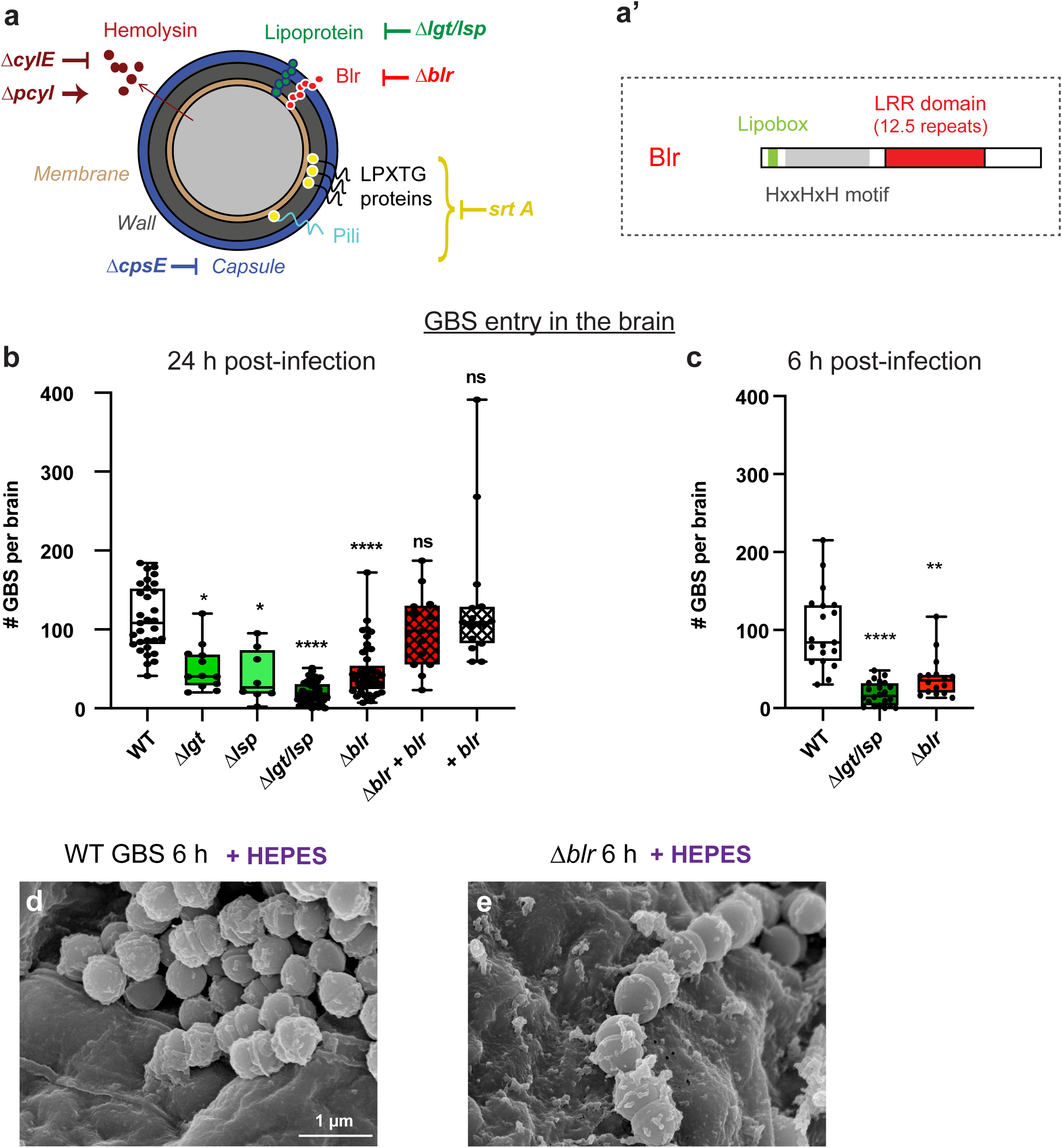
Screening for surface factors identifies the lipoprotein Blr as essential for BBB crossing in Drosophila. a-a’. Schematic representation of GBS surface structures and tested virulence factors with corresponding mutants. (A’). Schematic structure of Blr lipoprotein. b-c. Screening of GBS surface structures and virulence factors at (b) 24 h and (c) 6 h post-infection identified GBS surface lipoproteins, and in particular Blr, as crucial for BBB crossing. Kruskal-Wallis test. *(b) WT GBS (n = 31), Δlgt* (p = 0.0124, n=12), *Δlsp* (p = 0.0022, n = 8), *Δlgt/lsp* (*p* < 0.0001, n = 43), *Δblr* (*p* < 0.0001, n = 45), *Δblr*+*blr* (*p > 0*.*9999, n = 13), and* + *blr* (*p > 0*.*9999, n = 15). (c) WT GBS (n = 19), Δlgt/lsp* (*p <* 0.0001, n = 22), *Δblr* (*p* = 0.0029, n = 16). d-e. SEM picture of (d) WT GBS and (e)*Δblr* GBS attached to the brain surface.

Next, we sought to identify specific GBS lipoprotein(s) involved in BBB crossing. The GBS repertoire consists of 39 putative GBS lipoproteins^43^, most of them being substrate binding proteins of ATP-binding cassette (ABC) transporters. We selected Blr (group B leucine-rich), a His-triad/Leucine-Rich Repeat (LRR) protein, as an interesting candidate (Fig. 3a’). LRR domains are classically associated with protein-protein interaction and ligand recognition^44,45^. Similar LRRs are actually found within the internalin A^46^ (InlA) of *Listeria monocytogenes*, a surface protein crucial for bacterial crossing of the gut barrier^47^, albeit seemingly not of the BBB^48,49^. To test the role of *blr* (annotated as *gbs0918*), we deleted the gene in GBS NEM316. We first checked that GBS and its isogenic mutants grew similarly in various rich laboratory media (THY, BHI) at 37°C as well as in Drosophila Schneider medium at 30°C supplemented or not with HEPES (Suppl. Fig. 3b), ruling out any effect due to growth defect. Moreover, using SEM, we found no obvious difference in morphology between WT GBS and *Δblr*, that we found attached to the brain surface and in chains (Fig. 3d-e). However, as shown in Fig. 3b, *Δblr* displayed a significant decrease in bacterial count in the Drosophila larval brain at 24 h post infection compared to the control WT and complemented (*Δblr+blr*) strains. GBS translocation into the brain was also significantly decreased at 6 h post-infection for *Δblr* mutant (Fig. 3c). Altogether, these results showed that GBS lipoproteins, and in particular Blr, are key contributors to cross the Drosophila larval BBB and enter the brain *ex vivo*.

Surprisingly, we noticed that infection by Δ*blr* mutant resulted in significant damages to SPG membranes, altered septate junction architecture and increased BBB permeability compared to wild-type GBS (Supp. Fig. 2a-b and Supp. Fig. 3c). These differences were decreased but still remained when culture medium pH was maintained (Supp. Fig. 2a-b and Supp. Fig. 3c), and acidification was similar regardless of the bacterial strain (Supp. Fig. 2c), ruling out a differential effect from acidosis. Of note, none of these features were observed with the double Δ*lgt/lsp* mutant (Supp. Fig. 2a-b and Supp. Fig. 3c). In addition, using SEM, we detected the presence of additional, large structures reminiscent of the polysaccharidic coat produced during biofilm formation (Supp. Fig. 3d). Altogether, these data suggest that, in the absence of Blr, GBS turns on more destructive, yet much less efficient alternative mechanisms dependent on other lipoproteins. They also point to specific Blr-dependent mechanisms for GBS attraction to and crossing of the BBB.

### The Drosophila lipoprotein receptor LpR2 is essential in the BBB for brain invasion by GBS

We then asked how the lipoprotein Blr overcomes the physical barrier of the SPG (Fig. 1a-c). Interestingly, the LRR-containing InlA was shown to interact with human E-cadherin^50^. We tested the role of the Drosophila E-cadherin (*shotgun* gene, *shg*) in GBS entry *ex vivo*, and found that specifically knocking it down in the SPG layer, through the GAL4/UAS system^51^, did not affect GBS brain entry (Fig. 4a). This suggests that GBS does not rely on E-cadherin for entering the fly brain.

**Figure 4:**
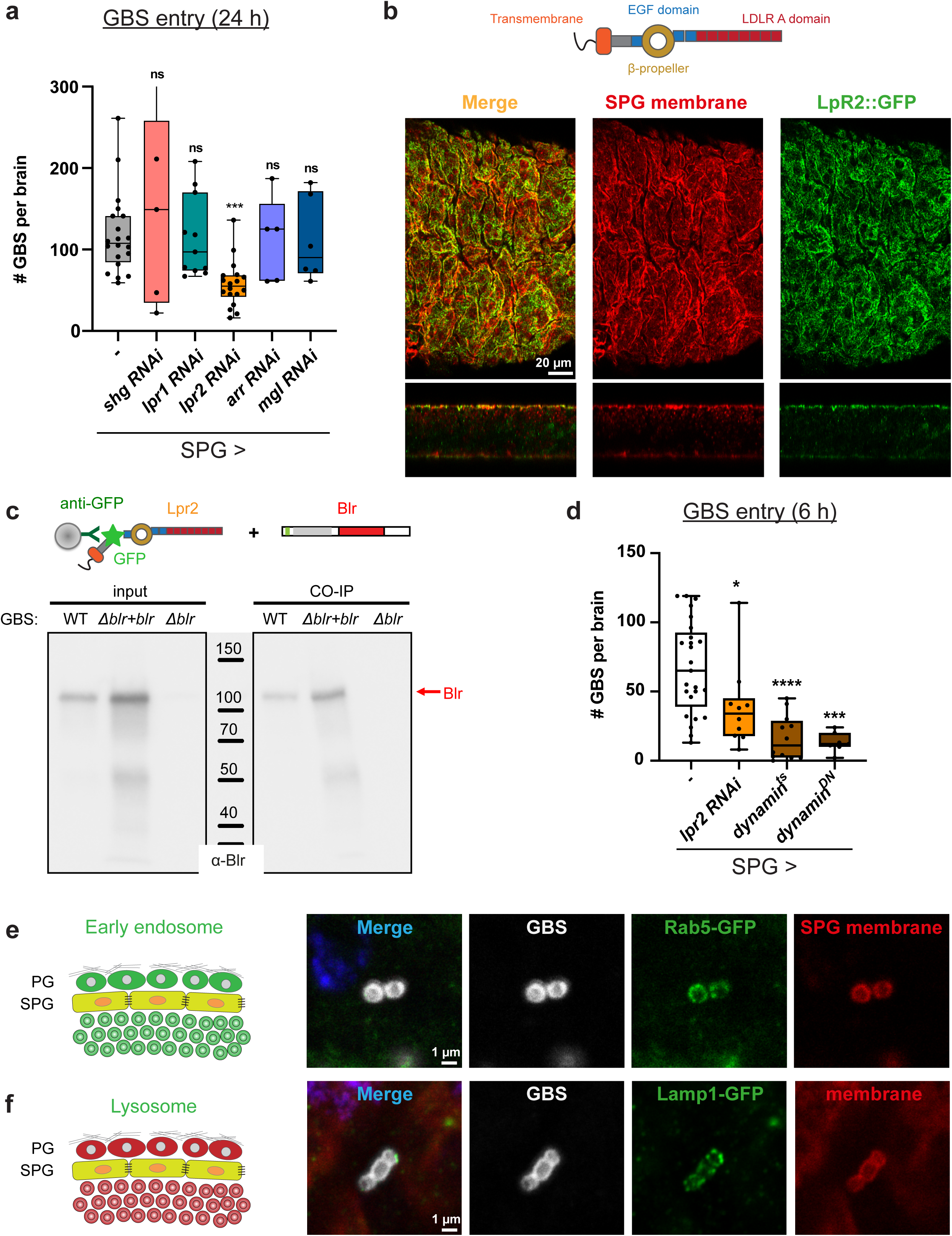
Drosophila lipoprotein receptor LpR2 mediates transcellular passage of the SPG by GBS through endocytosis. a. A knock-down screen for Drosophila E-cadherin (Shotgun) and lipoprotein receptors (LpR1, LpR2, Arrow and Megalin) identified LpR2 as crucial for BBB crossing by GBS. Kruskal-Wallis test of each knockdown versus control: control (n = 20); *shg*, p > 0.9999 (n = 5); *lpr1*, p > 0.9999 (n = 11); *lpr2, p = 0*.*0003 (n = 18); arr*, p > 0.9999 *(n = 5); mgl*, p > 0.9999 *(n = 6)*. b. Schematic representation of LpR2 structure and confocal image (top and orthogonal views) of LpR2::GFP genomic knock-in line (green) showing colocalisation of LpR2 on SPG membranes (*mdr65-mtd-Tomato*, red). c. Co-immunoprecipitation experiment between LpR2::GFP immobilised on beads and bacterial lysates of WT GBS, (*Δblr+blr*) GBS and *Δblr* GBS, detected with an antibody against Blr. A robust Blr-LpR2 interaction was revealed. d. GBS brain invasion is endocytosis-dependent. GBS entry at 6 h post infection was significantly decreased by either knocking down *lpr2* or blocking endocytosis (*dynamin*^*ts*^ and *dynamin*^*DN*^) specifically in the SPG. Mann-Whitney: p(control vs *SPG>lpr2 RNAi*) = 0.0162; p(control vs *SPG>dynamin*^*ts*^) < 0.0001; p(control vs *SPG>dynamin*^*DN*^) < 0.0001. Control (n = 25); *SPG>lpr2 RNAi* (n = 10); *SPG>dynamin*^*ts*^ (n = 12) and *SPG>dynamin*^*DN*^ (n = 7). e-f. Colocalisation of GBS (white) with markers for (e) early endosome (Rab5-GFP) and (f) lysosome (Lamp1-GFP) in green within the SPG membrane (*mdr65-mtd-Tomato*, red).

Lipoprotein receptors are conserved throughout the animal kingdom, and were originally identified as surface receptors capable of mediating cellular lipid uptake^52^. Lipids circulate in the blood in association with specific proteins called apolipoproteins (apo), forming lipoprotein particles of different densities (Low Density, LDL; Very Low density, VLDL). Lipoprotein receptors can be classified in two main groups, based on whether they behave as endocytic receptors supporting lipoprotein internalisation (LDLR, VLDLR, SR-A) or as mediators of lipid exchange at the cell surface. Of interest, Drosophila lipoproteins and lipoprotein receptors are similar to those in vertebrates^53,54^. Drosophila has seven lipoprotein receptors, belonging to the (V)LDLR families. Interestingly, specific lipoprotein particles were shown to cross the larval BBB, in which the receptors LRP1 and Megalin are expressed^30,55,56^.

We investigated which lipoprotein receptors were expressed in the SPG, and if they were important for GBS entry. Transcriptional data suggested expression of *lpr1, lpr2, arr and mgl* in the SPG (*Pauline Spéder and Andrea H. Brand, unpublished data*). Specific knockdown of these receptors identified LpR2 as the main lipoprotein receptor mediating GBS entry in the brain (Fig. 4a). More precisely, knocking down LpR2 in the SPG only was sufficient to decrease GBS count in the brain at 24 h post infection. In contrast, and in accordance with our previous findings, LpR2 was not required in the PG layer for GBS brain entry (Supp. Fig. 4a). Using a gene knock-in producing an endogenous LpR2-GFP fusion (LpR2::GFP MiMIC line), we observed that LpR2 expression was restricted to the SPG, where it colocalised with a membrane marker (Fig. 4b). Similar results were obtained using an anti-LpR2 antibody (Supp. Fig. 4b-c’’). These results show that LpR2, a lipoprotein receptor specifically expressed in the SPG, is crucial for GBS dissemination into the brain.

### GBS surface lipoprotein Blr binds to the Drosophila LpR2, allowing endocytosis-dependent transcellular crossing of the BBB

Interestingly, LpR2 has been shown to be an endocytic receptor, able to mediate the uptake of lipoprotein particles ^54,57^. We hypothesised that binding of LpR2 to Blr could first help GBS adheres to the BBB, and ultimately lead to its internalisation into the SPG through endocytosis. We first tested if LpR2 and Blr were able to physically interact. We set up a co-immunoprecipitation experiment between the two species, incubating bacterial lysate on LpR2-GFP fusions extracted from larval brains and bound to beads (see Methods). We showed that Blr was found in the bacterial eluates from LpR2-GFP beads for wild-type and complemented (Δ*blr+blr*) strains, whereas no band was recovered from Δ*blr* eluates (Fig. 4c and Supp. Fig. 4d). These data showed that Drosophila LpR2 is able to bind streptococcal Blr.

We further assessed the role of the endocytic pathway in GBS entry. We blocked endocytosis specifically in the SPG by preventing dynamin function through the overexpression of temperature-sensitive (*shibire*^*ts*^) or dominant-negative (*shibire*^*DN*^) forms. This led to a strong decrease in bacterial counts within the brain, detected as early as 6 h post-infection (Fig. 4d). In addition, we infected larval brains in which a marker for early endosomes (Rab5-GFP) was ubiquitously expressed along a specific marker for the SPG membrane. Under these conditions we were able to detect GBS in vesicles co-staining for Rab5-GFP and SPG membranes (Fig. 4e). Expressing another early endocytic marker (FYVE-GFP) specifically in the SPG gave similar results (Supp. Fig. 4e). Finally, we were able to detect GBS in lysosomal vesicles coming from the SPG layer, through the specific expression of the Lamp1-GFP marker (Fig. 4f). Taken together, these results strongly indicate that BBB crossing by GBS occurs via endocytosis, likely through binding of Blr to LpR2 and internalisation of the resulting complexes.

### Blr is a virulence factor essential for BBB crossing in the Drosophila larva

This brain explant protocol in the Drosophila larva led us to uncover and propose a novel mechanism for BBB crossing by GBS, in which the pathogen surface lipoprotein Blr interacts with the host lipoprotein receptor LpR2, leading to bacterial endocytosis and CNS invasion. To confirm these findings in an *in vivo* set-up, we developed a protocol of brain infection through pathogen microinjection into the Drosophila circulatory system (Fig. 5a). It was preferred to feeding in order to control the dose and bypass the variability in gut crossing efficiency.

**Figure 5:**
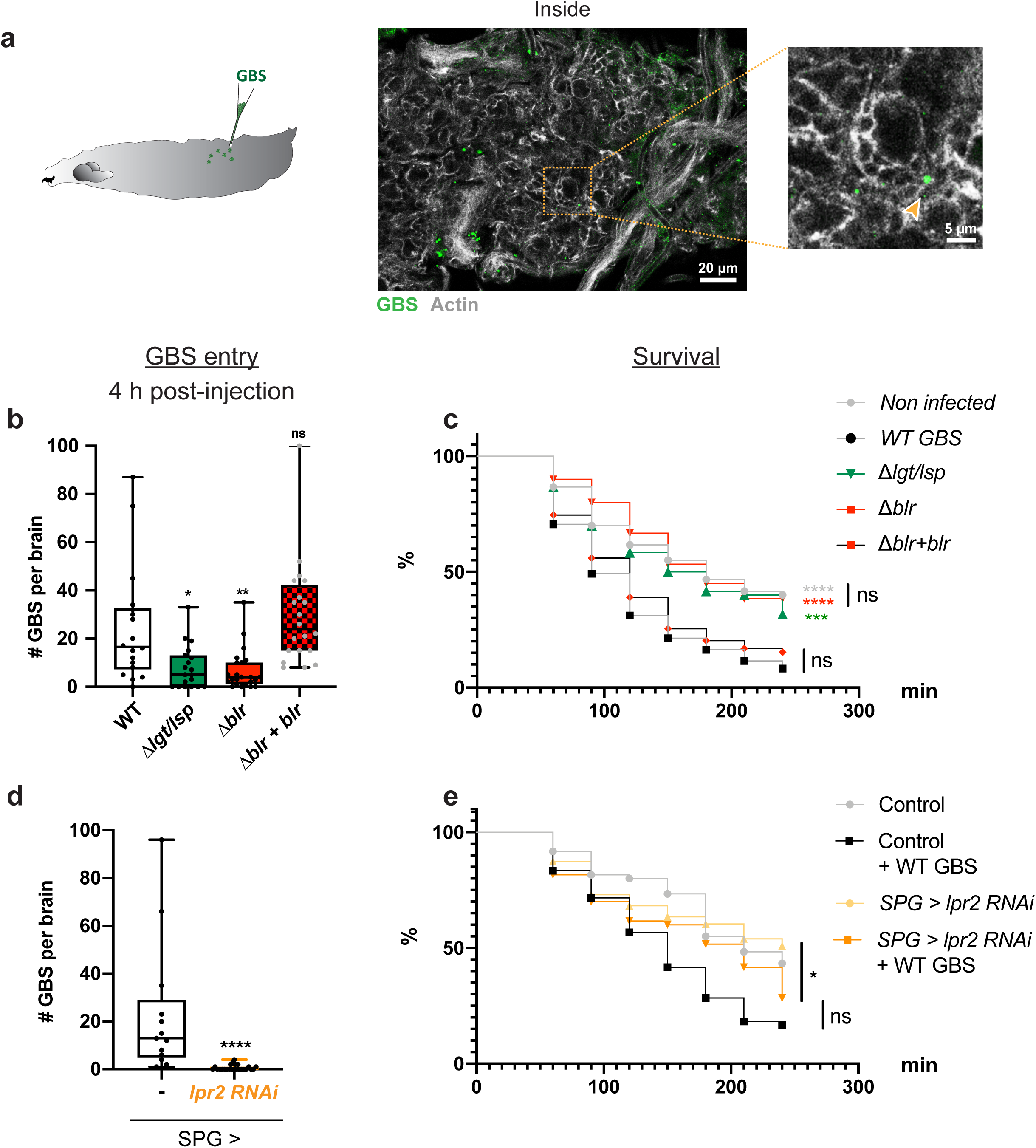
An *in vivo* model of brain infection in Drosophila identifies Blr as a virulence factor and confirms LpR2 as a BBB receptor for brain invasion by GBS. a. Schematic representation of Drosophila third instar larva injected with GBS. Confocal picture and close-up showing GBS (in green) inside the brain, 4 h after microinjection. b. GBS brain entry 4 h post-injection for WT GBS (n = 18), *Δlgt/lsp* (n = 19), *Δblr* (n = 23) and *Δblr+blr* (n = 20). Kruskal Wallis test: *Δlgt/lsp* p = 0.0168, *Δblr* p = 0.0039, *Δblr+blr* p *=* 0.6579. c. Kaplan-Meier survival curves for larvae injected with mock, WT GBS, *Δlgt/lsp, Δblr* and *Δblr+blr* strains (n = 60 for each condition) show that *Δlgt/lsp* and *Δblr* are less virulent than WT GBS and (*Δblr+blr*) GBS. Log rank test: p(GBS WT vs mock) < 0.0001; p(GBS WT vs *Δlgt/lsp*) = 0.0002; p(GBS WT vs *Δblr*) < 0.0001; p(GBS WT vs *Δblr+blr*) = 0.2797, and p(mock vs *Δblr*) = 0.9686. d. GBS brain entry at 4 h post injection in control (n = 13) and *lpr2* knockdown (n = 24) larvae. Mann Whitney test: p < 0.0001. e. Kaplan-Meier survival curves for control larvae and larvae in which *lpr2* has been knocked down in the SPG (*SPG>lpr2 RNAi*), injected with mock or WT GBS (n = 60 for each condition). Log rank test: p(Control + GBS vs *SPG>lpr2 RNAi* + GBS) = 0.0565; p(*SPG>lpr2 RNAi* vs *SPG>lpr2 RNAi* + GBS) = 0.0344.

Bacterial counts in the brain of surviving larvae at 4 h and 18 h post injection revealed that GBS was able to access and enter the Drosophila brain via the systemic route (Fig. 5b and Supp. Fig. 5a). Survival curves (0 - 4 h post injection) showed that systemic infection by wild-type GBS increased larval lethality compared to mock injection (Fig. 5c). These results demonstrated that GBS is able to infect the Drosophila brain from a circulating, systemic route, causing animal mortality.

Next, we tested the virulence of Δ*lgt/lsp* and Δ*blr* mutants in this set-up. First, bacterial counts in the brains of surviving larvae injected with Δ*lgt/lsp* or Δ*blr* were significantly reduced compared to WT or complemented (Δ*blr+blr*) GBS strains at 4 h post injection (Fig. 5b). To discard differences in fitness or survival between these isogenic GBS strains, we determined through cfu (colony-forming units) counts (see Methods) the exact quantity of bacteria per animal: in or attached to the brain, in the hemolymph, and in all other solid tissues (Supp. Fig. 5b). We then calculated three ratios: brain to hemolymph, brain to tissues, brain to hemolymph and tissues (Supp. Fig. 5c). In all cases, we found a significant decrease in Δ*blr* ratios versus wild-type ratios. This shows that the loss of Blr specifically affects the neurotropic ability of GBS to adhere and/or enter the brain. In agreement with these results, survival scores were significantly higher in larvae injected with Δ*lgt/lsp* or Δ*blr* mutants compared to the two control strains, with a lethality level similar to non-infected animals (Fig. 5c).

We then assessed the role of LpR2 in the BBB during systemic infection. Infection by WT GBS of larvae in which LpR2 was specifically depleted in the SPG resulted in a dramatic reduction of bacterial count in the brain (Fig. 5d). Survival curves showed that depleting LpR2 in the SPG led to a delayed lethality compared to wild-type animals (compare black and orange curves in Fig. 5e), although it did not reach statistical significance. This suggests that, although lethality might result from brain infection, it likely also depends on a systemic effect.

### Blr is a virulence factor essential for BBB crossing in mice

Combining *ex vivo* and *in vivo* infection protocols allowed us to propose Blr as a virulence factor of GBS essential for brain invasion in Drosophila. To determine whether this mechanism is conserved in mammals, we used the mouse model of GBS hematogenous brain infection^58^ and compared wild-type GBS strain with the isogenic *Δblr* mutant.

Time-course infection analysis showed that GBS could be detected in the brain as early as 3 h post-infection, was maintained at similar levels at 6 h and 24 h, and reduced at 48 h (Fig. 6a). In parallel, bacterial counts in the blood were measurable at 3 and 6 h post infection and dropped sharply at 24 h (Fig. 6b). Using a fluorescent GFP-tagged GBS, we observed bacteria attached to and in the capillaries of the brain parenchyma at 4 h post-infection (Fig. 6c, Supp. Fig. 6a) suggesting that the primary entry point for GBS is through the endothelial barrier. Interestingly, we were able to detect LDLR on these capillaries, underlying the availability of this receptor at GBS putative point of entry (Supp. Fig. 6b). Then, at 24 h after infection, we detected bacteria at the choroid plexuses and walls of the lateral (Fig. 6d), third, and fourth (not shown) ventricles, that also play a barrier role in the mammalian brain. Very few cells were detected in the brain parenchyma, in regions far from the ventricles, except for some small clusters in which typical streptococcal chains were identified (Fig. 6d).

**Figure 6:**
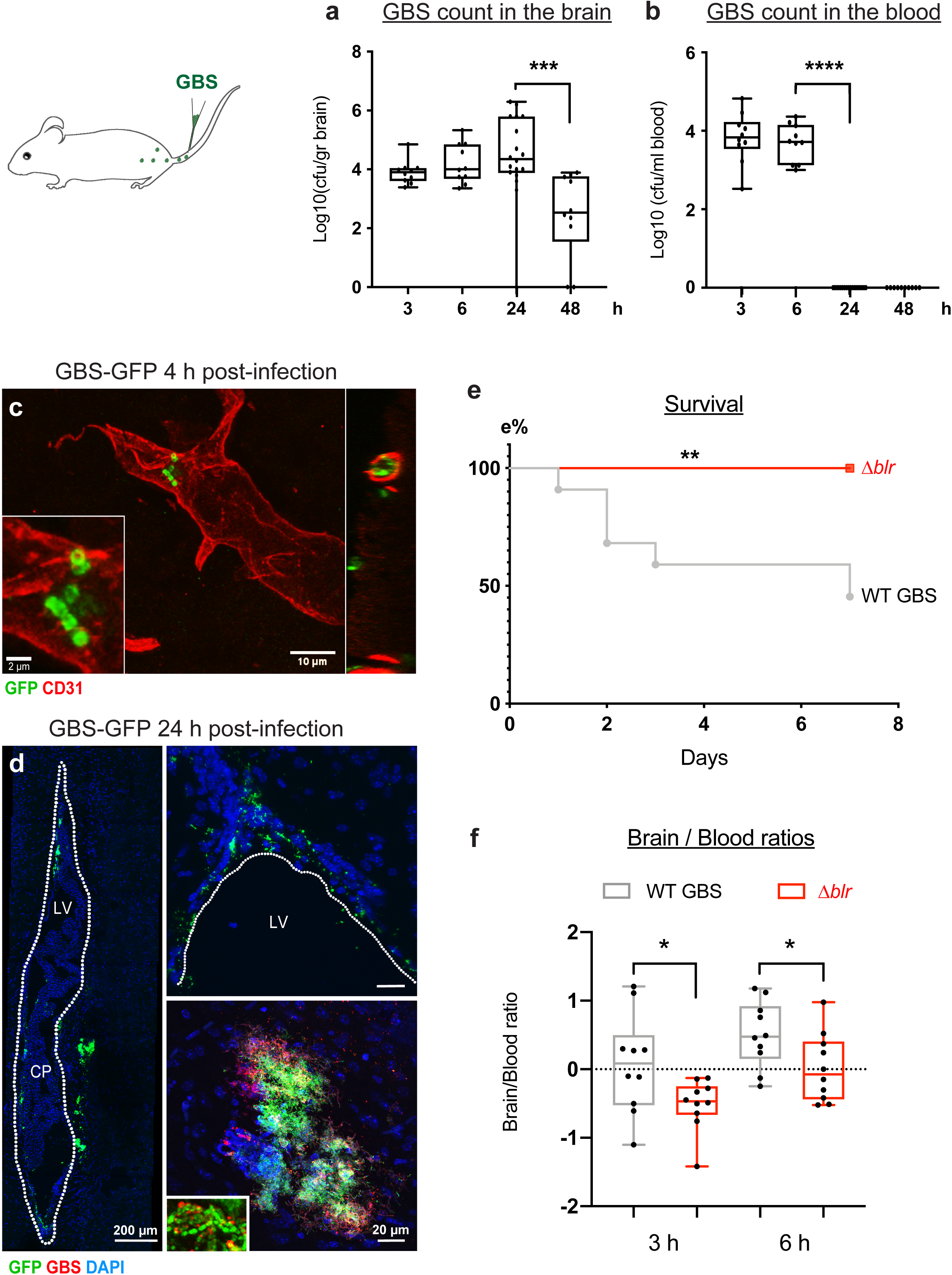
Blr is a streptococcal virulence factor in mice involved in BBB crossing by GBS. a-b. GBS counts in the (a) brain [log10(cfu/g)] and (b) blood [log10(cfu/ml)] of mice inoculated with GBS WT at 3 h (n = 10), 6 h (n = 10), 24 h (n = 18), and 48 h (n=10). Mann Whitney test: p(Brain 24 h vs 48 h) = 0.0002 and p(Blood 6 h vs 24 h) < 0.0001. c. Confocal images showing GBS-GFP (green) attached to and in the capillaries (CD31, red) of the brain parenchyma at 4 h post-injection. d. Confocal images of sagittal brain sections of mice injected with a fluorescent GBS WT-GFP strain demonstrating the presence of GFP-positive bacteria (green) at the choroid plexus (CP) inside the lateral ventricle (LV; outlined; left image) as well as at the walls of the LV and in the brain parenchyma adjacent to the LV (upper right image), at 24 h post infection. In the lower right image, a representative cluster of GFP-positive bacteria (also positive for anti-GBS; red) detected in the brain parenchyma. Typical streptococcal chains found in the clusters are presented in the inset. Nuclei were counterstained with DAPI and visualised in blue. e. Kaplan-Meier survival curves of mice intravenously injected with WT GBS (n = 22) or Δ*blr* (n = 10). Log-Rank test p = 0.0055. f. The ratio of bacterial counts in the brain versus blood [log10([cfu/g brain]/[cfu/ml blood])] in mice inoculated with *Δblr* was significantly lower than in mice inoculated with WT GBS, at 3 and 6 h post inoculation (n = 10 for each condition). Student’s t test, 3 h: p = 0.0351; 6 h: p = 0.0404.

Survival curves showed that infection with wild-type GBS led to more than 50% of lethality over 7 days (Fig. 6e). The mice that survived up to 7 days exhibited aberrant behavior indicative of neurological deficits, including unilateral palsy, immobilisation, and imbalance. Mood aberrations, such as isolation and lack of explorative behavior, were also observed. Moreover, the brains of these mice revealed meningitis hallmarks including meningeal thickening and leukocyte accumulation in the meninges compared with saline-injected control mice (Supp. Fig. 6c), as identified by co-staining for macrophages (CD68, pan-macrophage marker) and microglia (Iba-1, microglia/macrophage marker).

In contrast, no deaths were recorded in mice infected with *Δblr* mutant and their survival curve was significantly different compared to mice inoculated with WT GBS (Fig. 6e). We then analysed bacterial levels in the brain and in the blood over the course of infection. The levels of the Δ*blr* mutant in the blood were not significantly different from wild-type GBS neither at 3 h nor at 6 h post infection and we observed a similar clearance at 24 h (Supp. Fig. 6d). Importantly, the brain levels of the *Δblr* mutant at 3 h and at 6 h were lower, yet not significantly, and a significant reduction was then observed at 24 h post-infection when compared with the WT strain (Supp. Fig. 6e). Normalising brain-to-blood levels confirmed that the *Δblr* strain was significantly altered in its capacity to invade the mouse brain at 3 h and 6 h post infection, as compared to the wild-type (Fig. 6f).

Interestingly, none of the mice infected with the *Δlgt/lsp* mutant died, as observed with Δ*blr* mutants (Supp. Fig. 6f). Bacterial levels of *Δlgt*/*lsp* mutant were reduced both in the blood and brain compartments at 6 h post infection as compared to wild-type GBS (Supp. Fig. 6g). Yet, the brain-to-blood ratios were not significantly different between these two strains (Supp. Fig. 6h) suggesting that *Δlgt*/*lsp* mutants are generally less fit *in vivo*.

Altogether, these results identify Blr as a new, conserved virulence factor endowing GBS the ability to cross the BBB in Drosophila and mouse.

## Discussion

Here we propose an original model of brain infection, using the Drosophila larval brain, as a mean to investigate molecular and cellular mechanisms contributing to the crossing of the BBB. Amongst the various neurotropic pathogens tested in this model, Group B Streptococcus, a pathogen responsible for meningitis in neonates and immunocompromised adults, was the most efficient one and used herein. We were able to identify the surface lipoprotein Blr as a key contributor of specific GBS entry into the brain. Furthermore, we were able to show that Blr binds to LpR2, a host lipoprotein receptor expressed in the Drosophila BBB. We propose that binding of Blr to LpR2 promotes bacterial endocytosis, and GBS penetration into the brain through a transcellular passage (Fig. 7). Thus, using *in vivo* experiments in Drosophila, Blr and LpR2 were shown as key players in brain invasion. Blr contribution to CNS invasion was also established in a mouse model, demonstrating the power and relevance of the Drosophila model.

**Figure 7:**
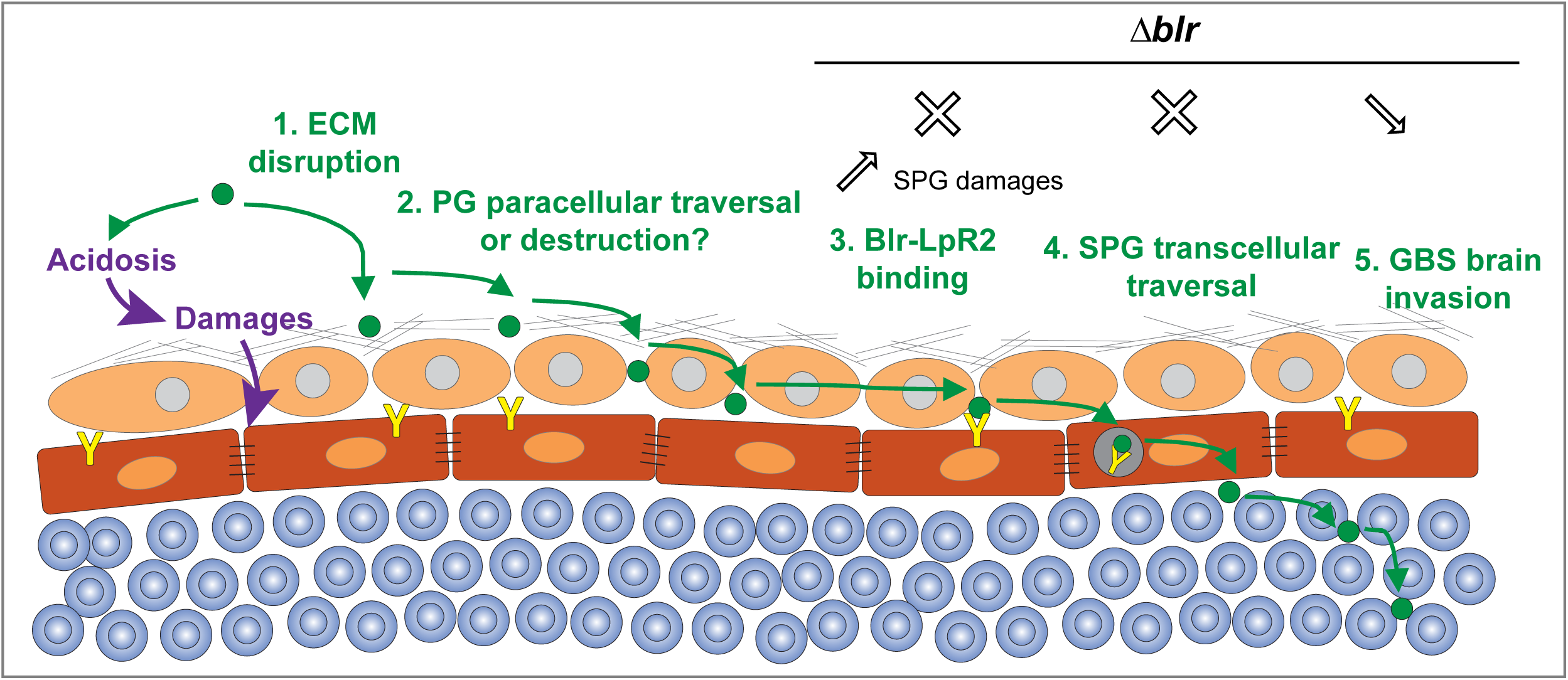
Proposed model for the mechanisms used by GBS during BBB crossing. GBS destroys the ECM layer and crosses the PG layer through a paracellular mechanism and/or cellular damages, likely supported or enhanced by acidosis. The bacteria adhere to the SPG surface via Blr-LpR2 interaction, allowing its internalisation through endocytosis and leading to its transcellular traversal. In the absence of Blr, GBS uses an alternative, albeit less efficient, mechanism for brain invasion, via SPG damages through an unknown process.

Our model combines an *ex vivo* approach with brain explants for straightforward, versatile and scalable screening of putative virulence factors and associated mechanisms, with a full *in vivo* approach to assess virulence and impact on the whole organism. Even though the *ex vivo* protocol does not allow to assess the contribution of circulating immune cells in BBB crossing, bypassing it can unveil BBB-specific mechanisms that could be masked either by an earlier, systemic effect (*e*.*g*., general inflammation) or by the difficulty to detect or assess it (*e*.*g*., acidosis). Interestingly, for example, *Cryptococcus neoformans* cannot enter the Drosophila larval brain in the *ex vivo* conditions (Fig. 1e), a finding congruent with the contribution of the Trojan horse mechanism proposed to explain *C. neoformans* barrier crossing ^59^.

Adult flies have been used previously as an infection model with GBS serotype Ia (A909 strain) through pricking, unveiling the role of GBS cell-wall anchored protein alpha C protein and host glycosaminoglycans as important for virulence and dissemination in the adult brain^60,61^. We chose to investigate the larval brain in order to study the impact of infection in a developing brain, where extensive neurogenesis is happening and neural circuits are being established, mimicking more closely neonatal infections caused by GBS.

Using our model, we demonstrated the key contribution of surface-exposed lipoproteins in mediating entry of GBS into the Drosophila larval brain, and in particular the role of a specific lipoprotein known as Blr. We focused our attention on Blr as an attractive candidate, because of its surface-exposed LRR region composed of 12.5 repeats of 22 amino acids. Blr was shown to be expressed *in vivo* but camouflaged by the bacterial capsule in GBS strain BM110. No role in virulence had been attributed to Blr yet. Indeed, Blr had been studied following intraperitoneal infection in a mouse model that bypasses the initial adhesion step. Interestingly though, it was shown that Blr can bind to the pathogen recognition receptor SR-A (scavenger receptor A), expressed on most macrophages and known to endocytose modified low-density lipoproteins^62^. This finding strongly supports our results that Blr interacts with a specific lipoprotein receptor LpR2 and is then internalised through endocytosis in the SPG. The physiological role of LpR2 in the BBB has not been described. In other tissues, LpR2, as well as the closely related LpR1, have been implicated in lipid metabolism^54,57^. Whole mutants for *lpr1* and *lpr2* are actually viable, albeit with lower fertility, suggesting that these receptors are not essential at least under homeostatic conditions.

During GBS infection, some bacterial lipoproteins are released in the extracellular environment and bind Toll-like receptor 2 through their lipid moiety^63^. However endogenous lipoprotein receptors, which are the natural receptor for lipoproteins *in vivo*, have been shown to bind lipoprotein complexes through their protein component (apolipoprotein)^64^. In addition, it is known that lipoprotein receptors bind most of their ligands through clusters of cysteine-rich LDL receptor type-A (LA) modules. LpR2, which bears between 7 and 9 LA motifs depending on the isoform^54^, could thus bind Blr through its protein moiety. The LRR domain (Fig. 3a’) would be of specific interest. Blr is also a virulence factor critical for BBB crossing in mice. LpR2 is orthologous to both LDLR and VLDLR proteins. Both LDLR and VLDLR have been shown to be expressed in brain endothelial cells, where they are linked to the uptake of molecular complexes across the BBB^65–67^. Here we confirmed LDLR localisation in blood vessels of the mouse brain *in situ* (Supp. Fig. 6b). Interestingly, VLDLR has been implicated in mediating the entry of Hepatitis C Virus into host cells^68^. The targeting of endogenous lipoprotein receptors could thus be a conserved strategy shared by a diversity of pathogens to enter cells and ultimately cross barriers, including the BBB.

Of note, we found that GBS acidifies the extracellular environment, a known parameter during meningitis. Although we demonstrated that it is not required for BBB crossing by GBS, it might help bacterial invasion, at least on a longer term, by weakening the SPG and/or affecting upstream layers, especially the PG. Indeed, lactic acid has been proposed as a virulence factor in rat fetal lung explants, where it is also linked to tissue destruction^69^. Interestingly, we noticed destroyed blood capillaries in the brain of mice infected with WT GBS (Supp. Fig. 6a), as well as brains with highly altered SPG during infection by WT GBS in our *in vivo* Drosophila model (data not shown). This suggests that acidosis-linked alteration of the BBB might be a conserved mechanism taking place during genuine infection.

Interestingly, Blr-deficient bacteria were less able to enter the brain while causing higher damages of the SPG. As mentioned previously, this suggests that when Blr is not available on their surface, bacteria turn to an alternative pathway, less efficient but more destructive, perhaps to gain access through alteration of junctions and tissue integrity. Such damages are not seen under infection with lipoprotein-deficient bacteria, in which GBS brain entry is extremely low. This indicates first that other lipoproteins support such a pathway, and also could explain why Δ*blr* still enters better than Δ*lgt/lsp*. The presence of biofilm is intriguing, and could be a way Δ*blr* causes additional damage to the BBB. Altogether, these different results underline the ability of GBS to shapeshift and use different mechanisms independently or together, depending on the conditions.

In conclusion, we propose the following model for GBS entry into the fly developing brain: rupture of the ECM, likely through GBS collagenolytic activity, and then traversal of the PG layer, through paracellular and/or destructive mechanisms. Then Blr comes at play, binds to LpR2 on the surface of the SPG allowing GBS endocytosis and brain invasion (Fig. 7). Our work, using an original model of brain infection in Drosophila, thus proposes a detailed mechanism behind pathogen crossing of the complex BBB structure, and identifies the specific lipoprotein Blr as a new, conserved virulence factor for GBS. Ultimately, understanding the exact cellular and molecular pathways employed by pathogens to use and corrupt the physiological role of host receptors will provide new insight into means of helping the BBB to restrict pathogen entry during infection as well as bypassing this barrier to target therapeutics molecules.

## Acknowledgments

We thank J. Culi, C. d’Enfert, G. Janbon, F. Leulier, M-K. Taha, M. Lecuit and F. Schweisguth for reagents and strains. We are very grateful to Gunnar Lindahl for the kind gift of his homemade antibody against Blr lipoprotein. L. Arbogast generated the *mdr65-mtdt-Tomato* construct. Stocks obtained from the Bloomington Drosophila Stock Center (NIH P40OD018537) and from the Vienna Drosophila Resource Center were used in this study. We are grateful to A-E. Deghmane, O. Disson, C. d’Enfert, G. Janbon, M-K. Taha and M. Lecuit for their kind and enthusiastic help and advice with the pathogen screen. We are indebted to D. Ferrandon for his expert advice on the project and to S. Liégeois for technical help, especially introducing us to hemolymph injection of Drosophila larvae. We thank B. Montagne for help with the Western blots. We thank E. Voulgari for valuable advice and S. Trygoni and D. Dionysopoulou for technical support on the mouse infection model. Further, we are grateful to the personnel of the Department of Animal Models for Biomedical Research of the Hellenic Pasteur Institute for their invaluable help. We thank D. Ferrandon for critical reading of the manuscript. This work has been funded by a starting package from Institut Pasteur/ LabEx Revive and a JCJC grant from Agence Nationale de la Recherche (NeuraSteNic, ANR-17-CE13-0010-01) to P.S.; a Grand Projet Fédérateur Microbes & Brains InFeSteR grant from Institut Pasteur to P.S., R.M. and S.D. B.B. has been supported by a Roux-Cantarini and a LabEx Revive post-doctoral fellowships.

## Author Contributions

B.B. performed most Drosophila experiments. N.R. generated Fig. 2A-B, Fig. 3C, Fig. 4B-C, Supp. Fig. 4D and parts of Fig. 4D and Supp. Fig. 2A, all under the supervision of B.B. P.S. devised the brain explant protocol and started the pathogen screen. B.B. and P.S. designed and analysed all Drosophila experiments. B.P. and S.D. generated GBS strains and constructs as well as performed growth curves. S.D. designed and advised on all GBS experiments. F.P. designed, performed and analysed, together with K.S., the mouse experiments and generated Fig. 6 and Supp. Fig. 6. V.M. provided consultation on the mouse infection model and resources. R.M. supervised and advised on all mouse experiments and analysis. C.S. generated the EM and SEM data. B.B., S.D. and P.S. wrote the article with input from FP and RM.

## Declaration of Interests

The authors declare no competing interest.

## Methods

### Animal models

#### Drosophila strains and larval culture conditions

The following fly stocks were used: wolbachia-free *w*^*1118*^ (used as reference strain for this work, ^70^), *mdr65-mtd-*tomato on (this study*), mdr65-Gal4* (BDSC 50472, ^71,72^), *UAS-mCD8-RFP* (BDSC 27399 and 27400*), NP6293-Gal4*; *tub-Gal80*^*ts*^ (Awasaki et al., 2008), *UAS-shg-RNAi* (BDSC stock 34831), *UAS-lpr-RNAi (VDRC stock 106364), UAS-lpr2-RNAi (VDRC stocks 107597 and 25684), UAS-arr-RNAi (VDRC stock 4818), UAS-mgl-RNAi (VDRC stock 105071), yw*; *Mi(PT-GFSTF*.*1)LpR2*^*MI04745-GFSTF*.*1*^*(BDSC stock 60219), UAS-shi*^*ts*^ (BDSC stock 44222), *UAS-shi*^*K44A*^ (BDSC stock 5811), *yw; EGFP-Rab5 (Fabrowski et al*., *2013), UAS-GFP-FYVE-myc (*BDSC stock 42716), *Vkg-GFP*^40^.

Embryos were collected for 2-3 h on grape juice egg-laying plates. Equivalent numbers (100) of hatching first instar larvae were transferred to standard food plates at 25°C or 29°C (for RNAi knockdown) until mid-third instar larval stage. For the *mdr65-Gal4, UAS-RFP x UAS-shi*^*ts*^, hatching first instar larvae were transferred to standard food plates at 18°C until early-third instar larval stage and transferred then to 30°C.

#### Microorganisms used and culture conditions

The microorganisms that were tested in our experimental set up are shown in Table 1. All strains were grown overnight at 37°C in BHI (Brain Heart Infusion) broth for bacteria or YPD (Yeast extract Peptone Dextrose) medium for fungi. They were stored at −80°C in BHI broth containing 20% glycerol for bacteria or in YPD broth containing 30% glycerol for yeast.

**Table 1.**
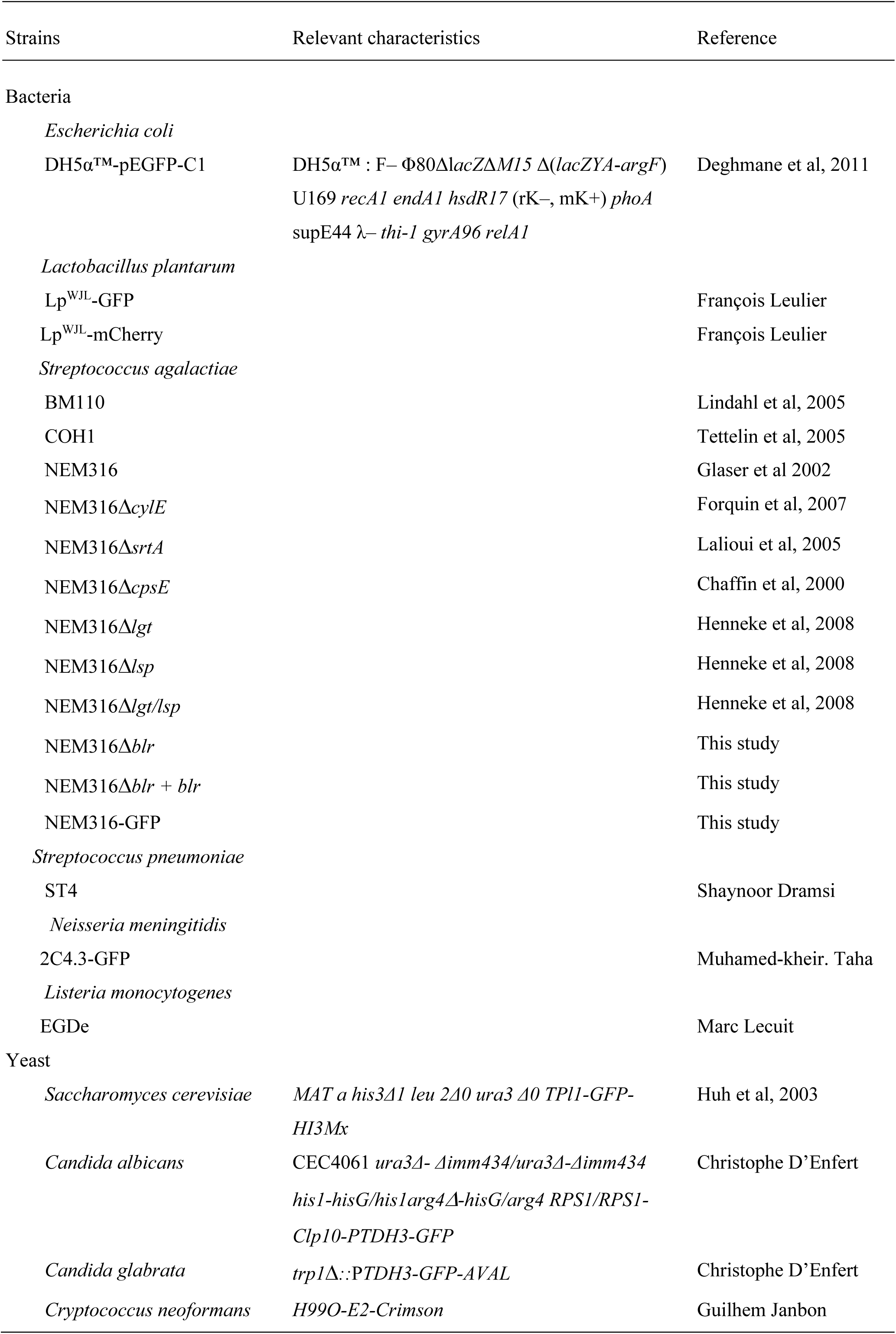
Microorganisms used in this study.

#### Mouse ethics statement

All animal experiments in this study were carried out in the Department of Animal Models for Biomedical Research of the Hellenic Pasteur Institute in strict compliance with the European and National Law for Laboratory Animals Use (Directive 2010/63/EU and Presidential Decree 156/2013), with the FELASA recommendations for euthanasia and Guide for the Care and Use of Laboratory Animals of the National Institutes of Health. All animal work was conducted according to protocols approved by the Institutional Animal Care and Use Committee of the Hellenic Pasteur Institute (Animal House Establishment Code: EL 25 BIO 013). *License No 6317/27-11-2017 for experimentation* was issued by the Greek authorities, i.e., the Veterinary Department of the Athens Prefecture. The preparation of this manuscript was done in compliance with ARRIVE (Animal Research: Reporting of *In Vivo* Experiments) guidelines.

## Protocols

### Construction of NEM316Δ*blr* mutant and complemented strain

In frame deletion mutant of *blr* in NEM316 was constructed by using splicing-by-overlap-extension PCR as previously described^73^. The primers used were the following:

- blr-1Eco 5’-TTCT*gaattc*TGTCGGTGCTGTAATGGAGT-3’

/ blr-2 5’-TAGCTCCGTAAAAGATTAGAGTCCTCCATAAATGT-3’

and

- blr-3 5’-AACATTTATGGAGGACTCTAATCTTTTACGGAGCTA-3’

/ blr-4Bam 5’-TTCT*ggatcc*AACCCCATGATGTAACACT-3’.

The chromosomal gene inactivation was carried out by cloning blr-1/blr-4 fragment into the thermosensitive shuttle plasmid pG+host5. Electroporation of the recombinant plasmid in *S. agalactiae* NEM316 strain and allelic exchange were performed as described^74^.

To complement the *blr* mutation in *trans*, the *blr* open reading frame was amplified using:

- pTCVblr-1Bam 5’-TCTC*ggatcc*TTATGGAGGACTCATGAAAG-3’ and
- pTCVblr-9BglII 5’-TCTC*gtcgac*GATTAATGGTGATGATGACC-3’ primer and cloned into the plasmid pTCV downstream from the constitutive promoter Ptet. The resulting plasmid pTCVΩPtet-*blr* was then transformed into competent NEM316Δ*blr* strain.

### Construction of GFP expressing NEM316

pMV158GFP is a mobilizable plasmid harboring the *gfp* gene cloned under the control of the P_M_ promoter^75^. pMV158GFP^Ery^ plasmid was constructed by replacing the Tc resistance gene of pMV158GFP by the *ermB* gene by using the Gibson method^76^. Briefly, *ermB* and pMV158-GFP were amplified with Erm-1 5’-GAGGGTGAAATATGAACAAAAA-3’ and Erm-2 5’-CCCTTAACGATTTATTTCCTCC-3’primers,andpMV158-35’-TTTTATATTTTTGTTCATATTTCACCCTCCAATAATGAGG-3’ and pMV158-4 5’-TATTTAACGGGAGGAAATAAATCGTTAAGGGATCAAC-3’,respectively. pMV158GFP and PCR product were ligated and the resulting pMV158GFP^Ery^ was used to transform *S. agalactiae* NEM316 strain, applying selection for erythromycin (10 µg/ml).

### Bacterial growth curves

One ml of overnight bacterial pre-culture in BHI was washed once in PBS and resuspended at OD600 of 2 ml^−1^. Then each culture was diluted in a given medium at 1/40 dilution and 180 µl of this suspension dispensed in 96 well plates in triplicate and absorbance measurements were recorded using a Biotek Synergy 2 microplate reader using Gen5 data analysis software (v.3.03).

### DNA cloning and Drosophila transgenics

A portion of the *mdr65* enhancer (GMR54C07, Flybase ID FBsf0000165529), which drives in the SPG, was amplified from genomic DNA extracted from *mdr65-GAL4* adult flies, with a minimal Drosophila synthetic core promoter [DSCP^71^] fused in C-terminal. The *mtd-Tomato* DNA codes for a Tomato fluorescent protein tagged at the N-terminal end with Tag:MyrPalm (MGCCFSKT, directing myristoylation and palmitoylation) and at the C-terminal with 3 Tag:HA epitope. It was amplified from genomic DNA extracted from *QUAS-mtd-Tomato* adult flies (BDSC30005, Chris Potter lab). The two amplicons were joined using the Multisite gateway system^77^ to generate a *mdr65*^*DSCP*^*-mtd-Tomato* construct. The construct was integrated in the fly genome at an attP2 docking site through PhiC31 integrase-mediated transgenesis (BestGene). Several independent transgenic lines were generated and tested, and one was kept (*mdr65-mtd-Tomato*).

### Culture of Drosophila brain explants

Staged larvae were washed successively in PBS and ethanol 70% v/v in water then transferred in cold Drosophila Schneider’s Medium in a dissection well. Larvae were cut at around a quarter from the posterior spiracle to minimise damages to motor nerves. The posterior part was discarded and the anterior part was turned inside out to expose the brain. All larval tissues were kept except for the gut, which is removed to avoid contamination with intestinal symbiotic pathogens. Eight larvae were transferred to one well (24-well cell culture plate: Falcon 353504) and cultured in 750 μl of Culture medium I (Drosophila Schneider’s medium (Gibco 217200-24) supplemented with 2 mM L-Glutamine (Gibco 25030-032) and 0.5mM Sodium L-ascorbate (Sigma A4034) at 30°C and 60% humidity under gentle rotary agitation (275 rpm on a Titramax 100 from Heidolph Instruments). After 3 h, the Culture medium I is replaced by Culture medium II [Culture medium I supplemented with 1% Fetal Bovine Serum (Sigma F4135)], then the medium was replaced after 3h and every 10h, by a fresh Culture medium II.

### Drosophila brain explants infection

An overnight preculture was set from glycerol stocks in BHI at 37°C for bacteria or in YPD at 30°C for yeast. The bacterial preculture was diluted 1/20 in BHI, and was grown for 2 h 30 min at 37°C (OD_600_ of 0.8). The yeast preculture was diluted to OD_600_=0.2 then grown 5 to 6 h at 30°C until OD_600_ of 1. A 10x infectious dose is then prepared after pelleting through 5 minutes centrifugation at 3,500Íg (at 4°C), washing each original culture twice in PBS, twice in Drosophila Schneider’s Medium and then resuspended in 750 µl of Schneider’s (10 × 10^8^ CFU/ml for *Streptococcus agalactiae, Streptococcus pneumoniae, Listeria innocua* and *Listeria monocytogenes*; 10 × 10^7^ CFU/ml for *Neisseria meningitidis, Candida albicans* and *Candida glabrata* and 10 × 10^5^ CFU/ml for *Cryptococcus neoformans*). Pathogen concentration was calculated by OD_600_ correlation (*Streptococcus agalactiae, Streptococcus pneumoniae, Listeria innocua* and *Listeria monocytogenes*: 1 OD_600_ = 8,8 × 10^8^ CFU/ml; *Neisseria meningitidis*: 1 OD_600_ = 10^9^ CFU/ml; *Candida albicans* and *Candida glabrata*: 1 OD_600_ = 3 × 10^7^ CFU/ml; *Cryptococcus neoformans*: 1 OD_600_ = 6 × 10^7^ CFU/ml).

The10 x infectious dose of each pathogen is diluted 1/10 in the brain explant culture medium I to reach the infectious dose (10^8^ CFU/ml). Brain explants were infected for 3 h at 30°C and 60% humidity under agitation (275 rpm on a Titramax 100 from Heidolph Instruments). Then, the infected medium was replaced by fresh culture medium II after 3 h and every 10 h.

### Dextran Permeability

We used 10 kDa Dextran (Texas Red, lysine fixable, D-1863, Invitrogen) at 50 mM final concentration in Culture medium II and we followed the same experimental and quantification procedures as described previously ^78^.

### DHE assay

To assess oxidative stress, we performed DHE (dihydroxyethidium) assay following standard procedures ^79^. Briefly, dissected brains were incubated for 5 min in 30 μM DHE, washed three times in PBS and then fixed for 8 minutes in 7% formaldehyde in PBS.

### *In vivo* Drosophila larval infection

GBS preculture and culture are prepared as described for the *ex vivo* protocol. 20 nl of concentrated GBS were injected in larvae using the nano-injector Nanoject III (Drummond Scientific) in order to reach 8.8 × 10^8^ CFU/ml of hemolymph.

### Immunohistochemistry

Brains were processed and stained according to standard procedures. Briefly, brains of inside-out larvae were fixed for 30 min in 4% methanol-free formaldehyde (Thermo Scientific, 28908) at room temperature, washed in PBS 3 × 10 min and permeabilised in PBS-Triton 0.3% for 3 Í 10 min. Brains were incubated with primary antibodies at 4°C in blocking solution (PBS-Triton 0.3%, Bovine Serum Albumin 5%, Normal Goat Serum 2%) for 18-36 h, then washed with PBS-Triton 0.3% and incubated with secondary antibodies 18-24 h at 4°C in blocking solution, and washed with PBS-Triton 0.3%, Samples were mounted in Mowiol mounting medium and visualised with a laser scanning confocal microscope (Zeiss LSM 880). The following primary antibodies or dyes were used: anti-GBS (homemade), anti-*S*.*pneumoniae* (homemade), anti-*L*.*monocytogenes* (R12, gift from M. Lecuit), anti-Blr^45^, anti-GFP (Abcam, ab13970), anti-LpR2 (gift from J. Culi), Phalloidin–Atto 647N (Sigma 65906), DAPI (Thermo 62247).

### Co-immunoprecipitation and western blot

For each condition, 100 brains of *yw; MiMIC(PT-GFSTF*.*1)LpR2*^*MI04745-GFSTF*.*1*^ larvae were dissected and lysed in lysis buffer (50mM Tris-HCl [pH 7.5], 150 mM NaCl, 1mM DTT, n-octyl-beta-glucopyranoside 1%, 5mM EDTA, 1mM PMSF, protease inhibitor cocktail Roche). Brain lysates were spun for 10 minutes at 4°C at 15,000Íg and incubated 1h at 4°C with 25 μl of equilibrated agarose beads (Chromotek, bab-20) to prevent non-specific binding to beads. The brain lysates were spun for 2 mins at 4°C at 2,500Íg and the cleared supernatant was incubated overnight at 4°C with 25 μl of equilibrated GFP-trap beads (Chromotek, gta-20). Bound GFP-trap beads were then washed three times, twice with lysis buffer and once with washing buffer.

Bacterial pellets of different GBS strains *WT, Δblr* and *complemented Δblr+blr* were lysed during 1 h at 4°C with 1 ml of lysis buffer. The bacterial lysates were spun for 10 min at 4°C at 15,000Íg and the supernatant was incubated for 1 h in 25 μl of equilibrated agarose beads at 4°C. The bacterial lysate was then spun for 2 min at 2,500Íg at 4°C and the cleared supernatant was incubated overnight at 4°C in the column containing bound GFP-trap beads. The column was spun for 2min at 2,500Íg at 4°C and the beads were washed three times, once with lysis buffer and twice with washing buffer. The beads were then resuspended in Laemmli 4X (BioRad) with 10% of β-mercapthoethanol and heated at 90°C for 10 min.

For western blot, proteins were boiled in Laemmli sample buffer, separated by SDS-PAGE on 7.5% Mini-Protean TGX Stain-Free precast Gels (Bio-Rad, 4568024), and transferred onto PVDF membrane using the Trans-Blot Turbo transfer pack (Bio-Rad). Immuno-detection was performed as follows: the membrane was blocked in PBS–skimmed milk 5% and incubated for 1 hour with rabbit primary anti-Blr (1/750) and mouse primary anti-GFP (Invitrogen, 1/1000, gift from B. Montagne) antibodies and then with the secondary Dylight_680_-coupled goat anti-rat antibody (Thermo Scientific Pierce) and HRP-coupled goat anti-rabbit antibody (1/10.000, gift from B. Montagne). Between the two antibodies and before detection, membranes were extensively washed with PBS + 0.1% Tween 20 and detection. was performed combining fluorescence and chemiluminescence (Bio-Rad ChemiDoc). For detection of brain lysate inputs, rat primary anti-GFP (1/1000, Chromotek [3H9]) and HRP-coupled goat anti rat antibody (S. Dramsi).

### Scanning Electron Microscopy (SEM)

Brains were fixed overnight in 2.5% glutaraldehyde in 0.1 M PHEM buffer pH 7.2. They were washed in 0.1 M PHEM buffer pH 7.2, post-fixed for 1 h and 30 min in 1% osmium tetroxide in 0.1 M PHEM buffer pH 7.2, and then rinsed with distilled water. Samples were dehydrated through a graded series of 25, 50, 75, 95 and 100% ethanol solutions followed by critical point drying with CO2.

Dried specimens were sputtered with 20 nm gold palladium, with a GATAN Ion Beam Coater and were examined and photographed with a JEOL JSM 6700F field emission scanning electron microscope operating at 7 Kv. Images were acquired with the upper SE detector (SEI).

### Transmission Electron Microscopy (TEM)

For transmission electron microscopy, brains were fixed with 2.5% glutaraldehyde in 1X PHEM buffer pH 7.2 overnight at 4°C. Specimens were post-fixed with tannic acid 1% in 0.1M PHEM buffer pH7.2 for 30’, post-fixed with 1% osmium tetroxide for 1 hr and 30 min in 0.1M PHEM buffer pH7.2 at room temperature, dehydrated in a graded series of ethanol, and embedded in Epon. After heat polymerization, thin sections were cut with a Leica Ultramicrotome Ultracut UC7’ sections (60 nm), stained with uranyl acetate and lead citrate. Images were taken with a Tecnai SPIRIT (FEI-Thermofisher Company at 120 kV accelerating voltage with a camera EAGLE 4K x 4K FEI-Thermofisher Company).

### Mouse infection

Eight to 10 week-old male CD-1 mice (body weight, 40.99 ± 3.62 g [mean ± standard deviation]) were randomly grouped and injected intravenously (i.v.), via the tail vein, with 10^8^ CFU of bacterial suspensions in sterile normal saline. For the determination of bacterial levels in blood and brain, mice were anaesthetized by intraperitoneal (i.p.) injection of a mixture containing ketamine (Imalgene 1000, MERIAL, Lyon, France; 100 mg/kg of body weight) and xylazine (Rompun, Bayer, Leverkusen, Germany; 10 mg/kg of body weight). Blood samples were collected by cardiac puncture. Immediately after, each mouse was euthanized by cervical dislocation and its brain was aseptically removed. One brain hemisphere from each mouse was homogenized in sterile normal saline. Bacterial levels in blood samples and brain homogenates were determined by plating serial tenfold dilutions on Columbia Agar with Sheep Blood plates (Thermo Fischer Scientific, Waltham, MA, USA) and counting of bacterial colonies 16 h later. The numbers of mice in each group of analysis are shown in Table 2. The bacterial loads per animal were then represented in a Log10 scale, and the Brain / Blood ratios were calculated as follows: Ratio Brain/ Blood = Log10 [(cfu / g brain)/(cfu / ml blood)]

**Table 2.** Sample size per time point per bacterial strain.

|  | Blood and brain levels |  |  |  | Survival |
| --- | --- | --- | --- | --- | --- |
|  | 3 h | 6 h | 24 h | 48 h | 7 d |
| WT GBS | 10 | 10 | 18 | 10 | 22 |
| *Δlgt/Δlsp* |  | 12 | 17 | 10 | 10 |
| *Δblr* | 10 | 10 | 9 |  | 10 |

### Mouse immunohistology

Mice were euthanized by (i.p.) injection of a ketamine/xylazine mix. After transcardial perfusion with 4% paraformaldehyde in phosphate-buffered saline (PBS), the brains of the infected mice were dissected out, post-fixed in the same fixative, cryoprotected in 30% w/v sucrose solution in PBS for 2 d at 4 °C, embedded in O.C.T. compound (VWR Chemicals) and frozen at −80 °C. Series of coronal or sagittal 20 μm-thick sections were collected on Superfrost Plus microscope slides and stored at −20 °C until further processing. The cryosections were thawed and subjected to antigen retrieval in 10 mM sodium citrate solution, pH 6, followed by 1 h blocking of non-specific sites with 5% v/v normal donkey serum (NDS), simultaneously with permeabilization using 0.1% v/v Triton X-100 in PBS. Primary antibodies diluted in 2.5% NDS in PBS were applied overnight at 4 °C, followed by incubation with the appropriate secondary antibodies for 2 h at room temperature. The following primary antibodies were used: rat anti-Cluster of Differentiation 68 (CD68; 1:100; Bio-Rad Antibodies, Oxford, UK; MCA1957GA), rabbit polyclonal anti-ionized calcium-binding adapter molecule 1 (Iba-1; 1:400; FUJIFILM Wako Pure Chemical Corporation, Osaka, Japan; 019-19741), rabbit anti-CD31(1:50; Abcam, Cambridge, UK; ab28364), goat anti-LDLR (1:100, R&D Systems, MN, USA; AF2255), rabbit anti-GBS (1:300; home-made). Secondary antibodies (all from Thermo Fisher Scientific) used for immunofluorescence were conjugated with Alexa Fluor 488 or 546 and cell nuclei were counterstained with 4′,6-diamidino-2-phenylindole (DAPI; 1:1000; Thermo Fisher Scientific). Prolong Gold antifade curing mountant (Cell Signaling Technology, Danvers, MA, USA) was used for mounting. Images were acquired using Leica TCS SP8 confocal microscope.

### Image processing

Fiji, Icy or Volocity were used to process confocal data. Adobe Photoshop and Illustrator were used to assemble figures.

## Statistics

GraphPad Prism software was used for all analyses.

### Bacterial quantifications in infected Drosophila brain

The same region of the CNS (Ventral Nerve Cord, VNC) was scanned at an optimised number of slices using a Zeiss LSM880 microscope. The exact number of bacteria for each brain was then determined manually by counting each individual bacterium contained within the boundary of the BBB (*mdr65-mtd-Tomato*).

### CFU counts following *in vivo* larval GBS-injection

Each injected larva is washed on a paper with ethanol 70% then bled in 10 µl PBS. The brain is then dissected, transferred and homogenized in 10 µl PBS. The rest of the larval carcass (other tissues except the gut) is also transferred and homogenized in 10 µl PBS. This protocol was done for 5 larvae by condition. Hemolymph, brain and carcass bacterial levels were determined by plating 7 serial tenfold dilutions two times on Columbia Agar with Sheep Blood plates (Biomérieux 43041) and counting of bacterial colonies after 16 hours at 37°C. The average CFU/µl was calculated as an average from all the different dilutions. The bacterial loads per animal were then represented in a Log10 scale, and ratios were calculated from raw counting then represented in a Log10 scale:

- Ratio Brain / Hemolymph = Log10 (cfu per brain / cfu per hemolymph)
- Ratio Brain / Other tissues = Log10 (cfu per brain / cfu per Other tissues)
- Ratio Brain / (Hemolymph+Other tisses) = Log10 (cfu per brain / (cfu per hemolymph+ cfu per Other tissues))

#### Drosophila statistical analysis

- In order to perform statistical tests on several experimental replicates, each value (corresponding to one brain) was normalised to the mean of the control condition within one replicate. Statistical tests were then run on all the normalised values from all replicates, which were considered as biological replicates.
- Comparisons between BBB permeability, GBS entry into the brain, cell viability, oxidative stress, bacterial levels in the hemolymph, bacterial levels in other tissues, bacterial levels in the brain, ratio of bacterial levels for brain/hemolymph, ratio of bacterial levels for brain/other tissues and ratio of bacterial levels for brain/other tissues+hemolymph were performed by Student’s t test (2 conditions) or one-way ANOVA test followed by Tukey’s post-hoc analysis (more than 2 conditions) when values followed a normal distribution (assessed by Shapiro-Wilk normality test). Otherwise, non-parametric Mann-Whitney tests (2 conditions) or Kruskal-Wallis tests (more than 2 conditions) were performed.
- Comparison of survival curves was performed using Log-rank test.
- P values lower than 0.05 were considered significant.

#### Mouse statistical analysis

- Comparisons between bacterial levels in the blood and the brain, as well as between ratios of bacterial levels for brain/blood were performed by unpaired Student’s t test when values followed a normal distribution (assessed by D’Agostino-Pearson normality test). Otherwise, non-parametric Mann-Whitney test was performed.
- Comparison of survival curves was performed using Log-rank test.
- P values lower than 0.05 were considered significant.

