## Supplementary materials for "An original model of brain infection identifies the hijacking of host lipoprotein import as a bacterial strategy for blood-brain barrier crossing"

#### Supplementary figure legends

**Figure S1: Drosophila as a platform to screen for mammalian neuro-invasive pathogens. Related to Figure 1.**

a. Cell death monitored by DAPI penetration in freshly dissected (n = 6), after 24 h of *ex vivo* culture (n = 8), and fixed (n = 2) brains. Cell viability is not affected in *ex vivo* compared to freshly dissected brains. Mann-Whitney test: p(fresh vs *ex vivo*) = 0.1079.

b. BBB permeability after 24 h *ex vivo* culture showed no significant effects. Mann Whitney test: freshly dissected (n = 11) vs *ex vivo* (n = 16) ns, p = 0.0989.

c. Oxidative stress monitored through DHE (Dihydroethidium) staining in brains freshly dissected (n = 7), after 24h of *ex vivo* culture (n = 7), or exposed to 0.1% (n = 5) $H_2O_2$. ANOVA test: p(fresh vs *ex vivo*) = 0.2096, p(*ex vivo* vs *ex vivo*+0.1%$H_2O_2$) = 0.0172.

d. Representative confocal images of Drosophila larval brains stained for phospho-histone 3 (PH3, red) after 6, 18, and 24 h of *ex vivo* culture.

**Figure S2: GBS uses a variety of mechanism to cross cellular layers. Related to Figure 2.**

a. Confocal images of the Drosophila BBB (top view and 3D orthogonal view) labelled for the SPG membrane in red (*mdr65-mtd-tomato*) at 6 h post-infection with WT, *Δblr* and *Δlgt/lsp* GBS strains (all green). Higher alteration of SPG labelling was observed under infection by *Δblr* GBS compared to infection by WT and *Δlgt/lsp* GBS. Such alteration was partially rescued by buffering medium acidity.

b. Close-up of confocal images of non-infected brain, brain infected with WT GBS and brain infected with *Δblr*, with and without acidosis, showing the SPG membrane (*mdr65-mtd-tomato,* red) and septate junctions (*Lachesin::GFP*, green) at 6 h post infection. Septate junctions are strongly affected under *Δblr* infection without HEPES and partially rescued with HEPES. SPG membranes are still damaged under *Δblr* infection with HEPES (6 h post-infection).

c. pH measurements of culture media non-inoculated and inoculated with WT, *Δlgt/lsp,* and *Δblr* GBS and *L.plantarum* for 3 h.

d-e. Transmission electron microscopy (TEM) pictures of a septate junction in (d) a control brain without infection and (e) in a brain infected 6 h under HEPES by WT GBS.

f. Scanning electron microscopy (SEM) picture of WT GBS bacteria attached to the surface of a Drosophila larval brain, under HEPES at 6 h post-infection.

**Figure S3: Roles of selected GBS factors on pathogenic invasion, BBB parameters and bacterial fitness. Related to Figure 3.**

a. Screening of selected GBS virulence factors and surface molecules at 24 h post infection identified GBS lipoproteins (*Δlgt/lsp* mutant*)* as a crucial factor for GBS brain invasion. Kruskal-Wallis test: WT GBS (n = 31); *cyl+* (n = 16), p = 0.0239; *ΔcylE* (n = 16), p > 0.9999; *ΔSrtA* (n = 13), p > 0.9999; *ΔcpsE* (n = 19), p > 0.9999; *Δlgt/lsp* (n = 43), p < 0.0001.

b. Growth curves of WT, *Δlgt/lsp, Δblr*, and *Δblr+blr* GBS strains at 30 °C in Drosophila Schneider's medium complemented or not with HEPES, and at 37 °C in THY and BHI media.

c. BBB permeability tests for brains infected by the different GBS strains (6 h post-infection) with and without HEPES. In both cases, BBB permeability is higher in brains infected with GBS-*Δblr* compared to other strains. ANOVA tests. Without HEPES: p(WT GBS vs *Δlgt/lsp*) = 0.1036; p(WT vs *Δblr*) = 0.0002. WT GBS (n = 7); *Δlgt/lsp* GBS (n = 7) and *Δblr* GBS (n = 8). With HEPES, ANOVA: p(WT vs *Δlgt/lsp*) = 0.9923; p(WT vs *Δblr*) < 0.0001. WT GBS (n = 8), *Δlgt/lsp* GBS (n = 7) and *Δblr* GBS (n = 8). p(*Δblr* vs *Δblr*+HEPES) < 0.0001.

d. Close-ups showing putative biofilms on brains infected by *Δblr*.

**Figure S4: The endocytic receptor LpR2 mediates specific passage of the SPG by GBS. Related to Figure 4.**

a. GBS brain entry is not affected, 24 h post-infection, after knocking down E-cadherin (*shg*) and lipoprotein receptors in the PG of Drosophila. ANOVA test: - (n = 14); E-cad, p = 0.9885 (n = 8); LpR1, p = 0.9858 (n = 7); LpR2, p = 0.4932 (n = 6); arr, p = 0.9886 (n = 6); Mgl, p = 0.8845 (n = 7).

b-c". Confocal images of (b-b") top close-up view and (c-c") orthogonal view showing LpR2 (anti-LpR2, white) and SPG (*mdr65-mtd-tomato,* red) colocalisation in the brain of a third instar larva.

d. Brain lysate input and unbound LpR2::GFP from the three co-immunoprecipitation columns.

e. Colocalisation of GBS (white) with both endosomal and lysosomal markers (*FYVE-GFP* and *Spin-RFP*, respectively in green and red), pinpointing GBS presence in autophagosomes.

**Figure S5: GBS brain entry *in vivo* through pathogen microinjection into the larval hemolymph. Related to Figure 5.**

a. GBS brain entry at 16 h post injection (lower inoculum of $10^5$ GBS/ml of hemolymph) for WT (n = 14) versus *Δlgt/lsp* (n = 24) and *Δblr* (n = 6) GBS. Mann-Whitney test: p(*Δlgt/lsp* vs WT) = 0.0011, and p(*Δblr* vs WT) = 0.0393.

b. Cfu counts in the brain, hemolymph and other tissues at 4 h after injection in the hemolymph of WT larvae of WT GBS (n = 5) and *Δblr* GBS (n = 5). Student's t test for each compartment: p(brain) = 0.0157, p(hemolymph) = 0.0341, p(other tissues) = 0.0562, p(other tissues+hemolymph) = 0.0063.

c. Ratio (in Log10) of bacterial counts in the brain versus hemolymph, brain versus other tissues, and brain versus other tissues+hemolymph at 4 h post injection.

Student's t test for each compartment: p(brain/hemolymph) = 0.0010, p(brain/other tissues) = 0.0064, p(brain/other tissues+hemolymph) = 0.0011.

**Figure S6: GBS infection in mice. Related to Figure 6.**

a. Confocal images showing GBS (green) around destroyed brain capillaries (CD31, red) at 4 h post-infection.

b. Confocal image of a sagittal mouse brain section displaying colocalisation of LDLR (green) and CD31 (endothelial cell marker; red) on brain capillaries.

c. Confocal images of sagittal brain sections of mice injected with saline or GBS WT at 7 d after inoculation, immunostained against Cluster of Differentiation 68 (CD68; green) and ionized calcium-binding adapter molecule 1 (Iba-1; red). Meningitis hallmarks including meningeal thickening and leukocyte accumulation in the meninges are apparent in the case of GBS WT-inoculated mouse as compared with the saline-injected control.

d. Blood bacterial levels [log10(cfu/ml)] for mice infected with WT and *Δblr* GBS at 3 (n = 10 for both), 6 (n = 10 for both), or 24 h (n = 18 and n = 17, respectively) after inoculation.

e. Brain bacterial levels [log10(cfu/g)] for GBS WT and GBS *Δblr* infected mice at 3 (n = 10 for both), 6 (n = 10 for both), or 24 h (n = 18 and n = 17, respectively) after inoculation. Mann-Whitney test: p(24 h) = 0.0009.

f. Kaplan-Meier survival curves of mice infected with WT (n = 22) or Δ*lgt/lsp* (n = 10) GBS. Log-Rank test: p = 0.0055.

g. GBS levels in the brain [log10(cfu/g)] and blood [log10(cfu/mL)] at 6 h post-injection for mice infected with WT (n = 10) and *Δlgt/lsp* (n = 12) GBS. Student's t test: p(brain) = 0.0009, and p(Blood) = 0.0023.

h. Brain/Blood ratio at 6 h post-injection. Student's t test*:* p = 0,6085.

### Supplementary figure 1

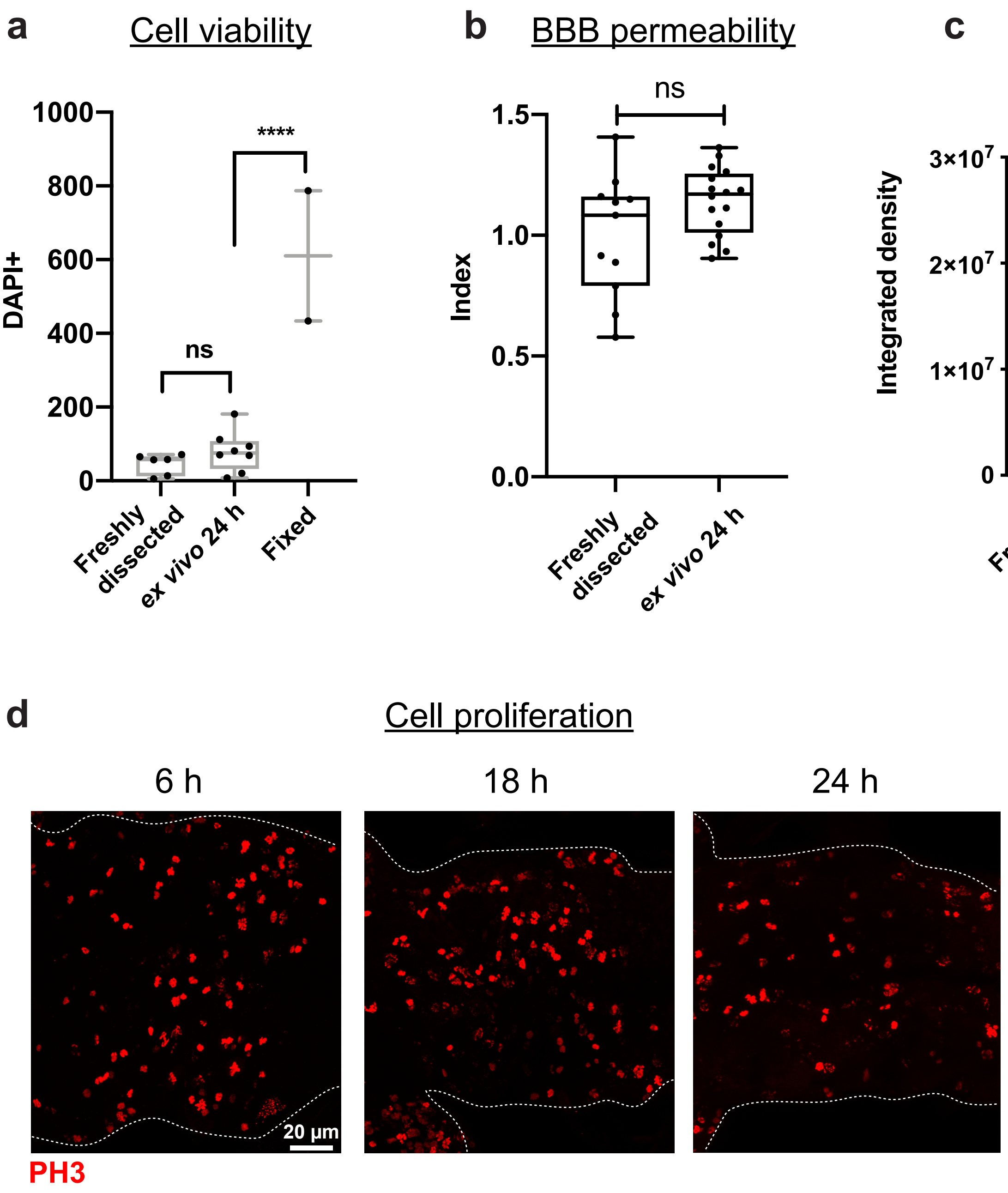

### Supplementary figure 2

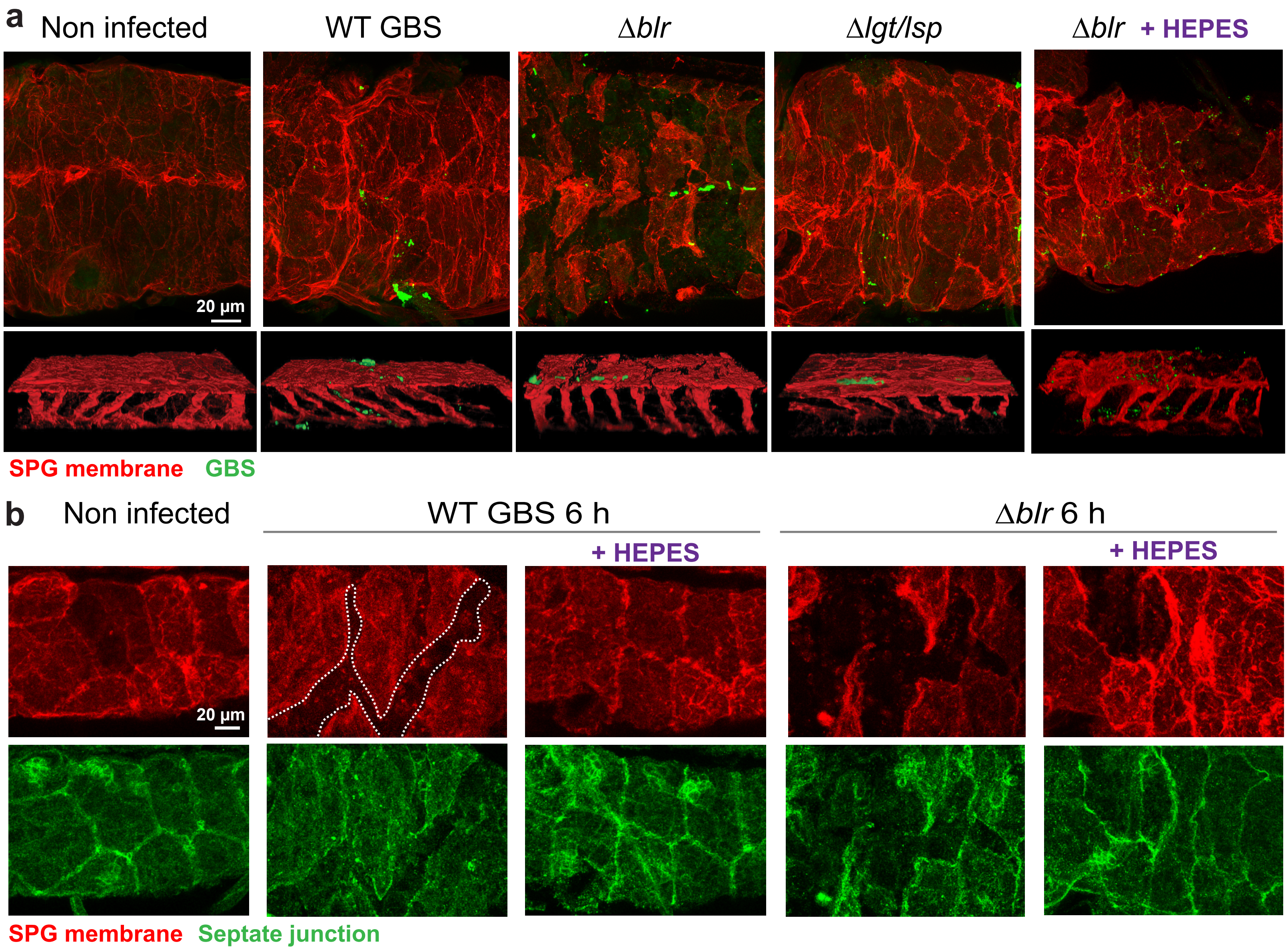


**c**

| | | 3 h infection |
|---|---|---|
| | - | 7 |
| GBS | WT | 4,6 |
| | *Δlgt/lsp* | 4,6 |
| | *Δblr* | 4,6 |
| | *L. plantarum* | 6,2 |

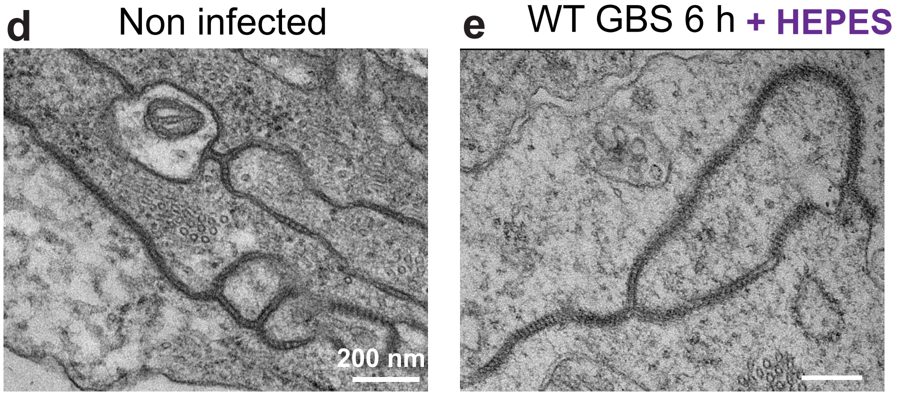


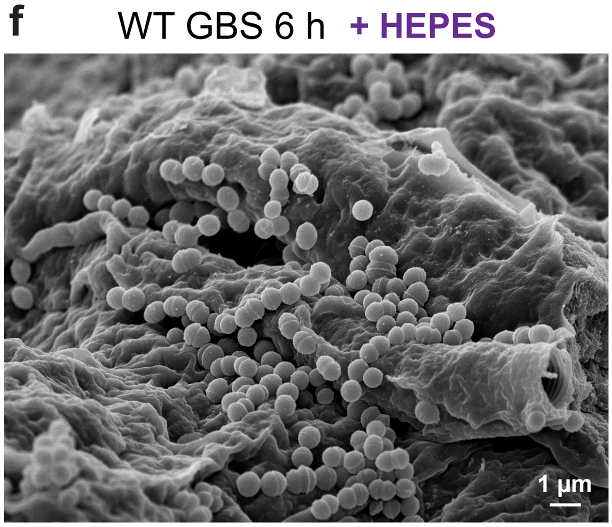

### Supplementary figure 3

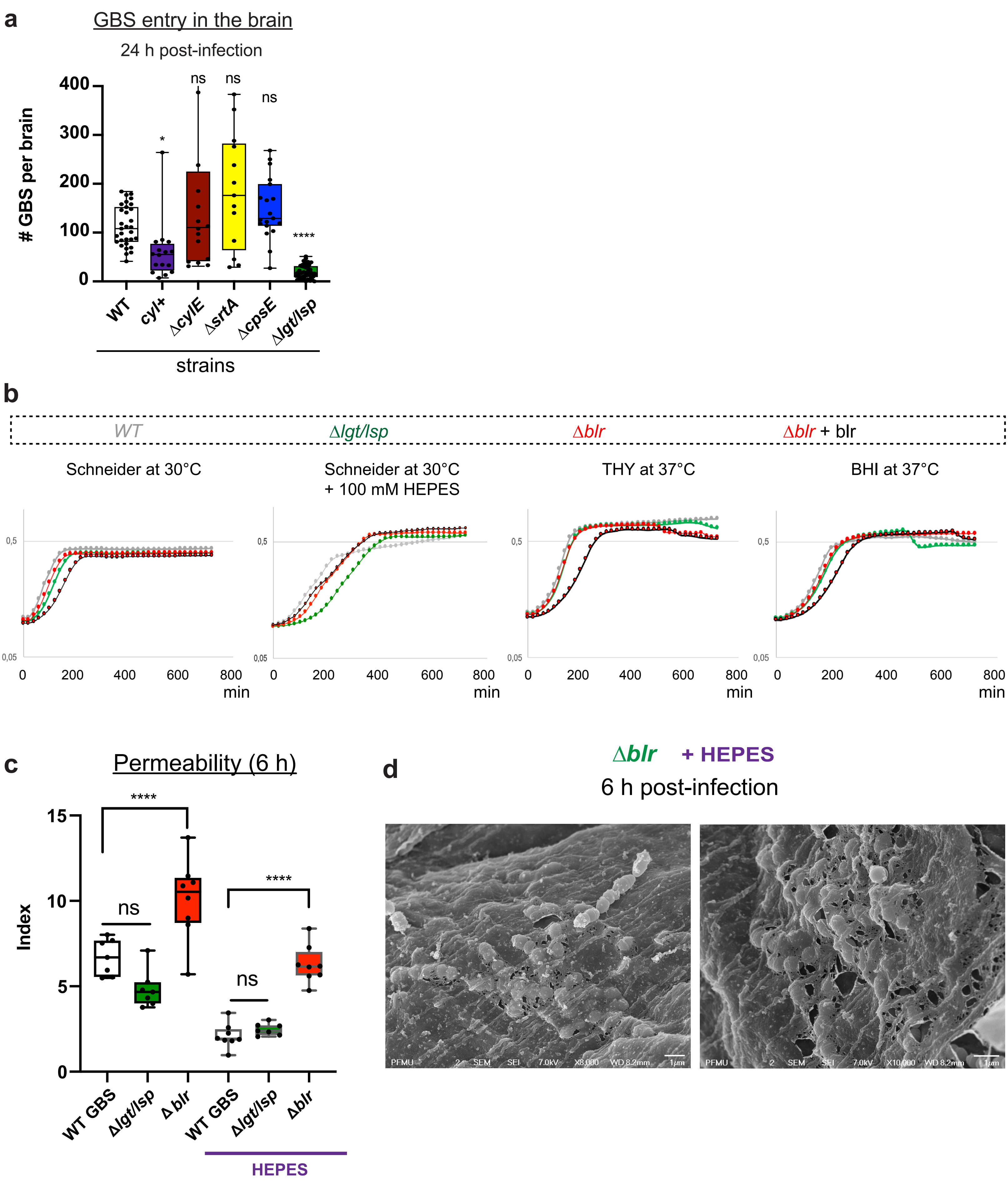

### Supplementary figure 4

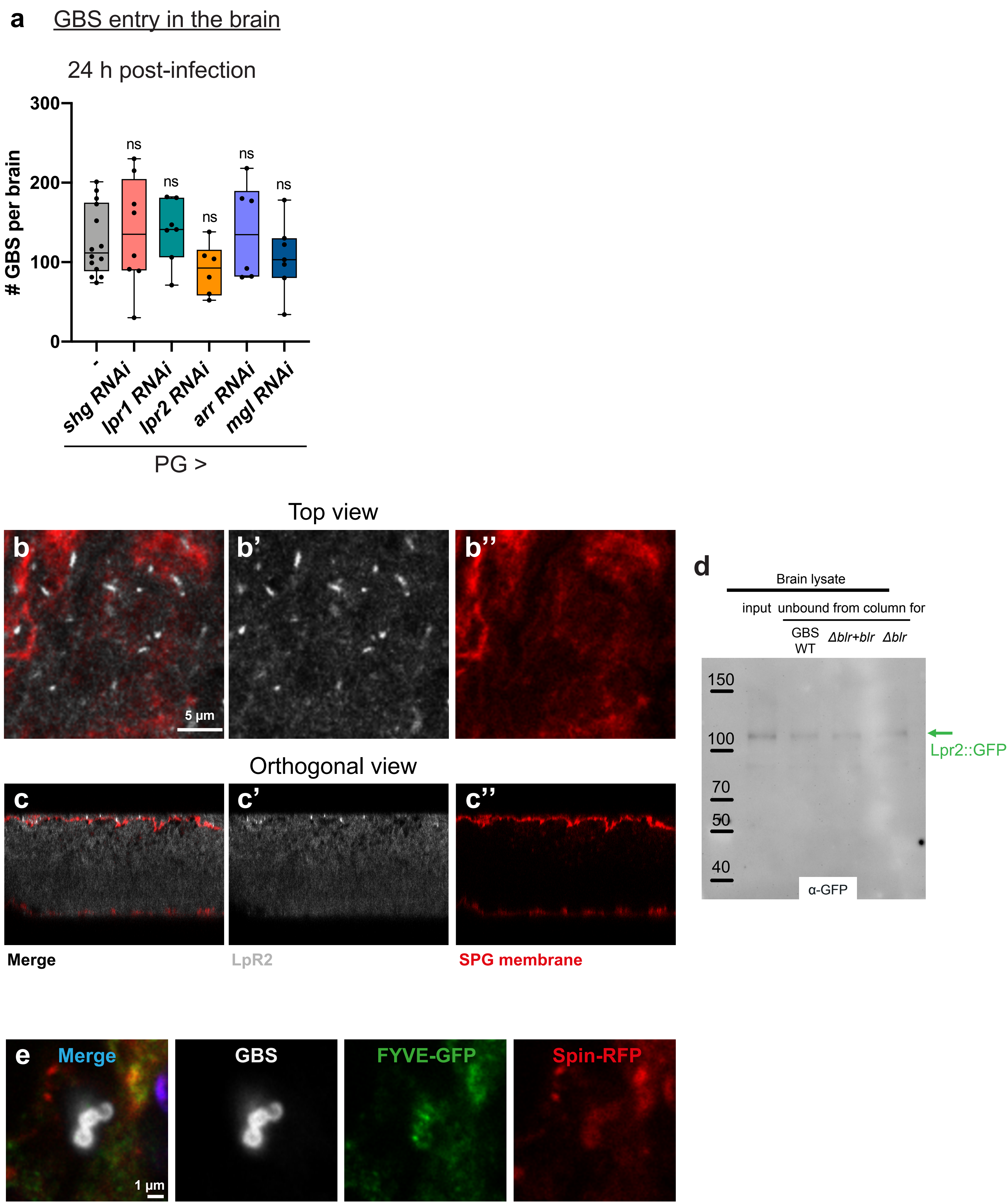

### Supplementary figure 5

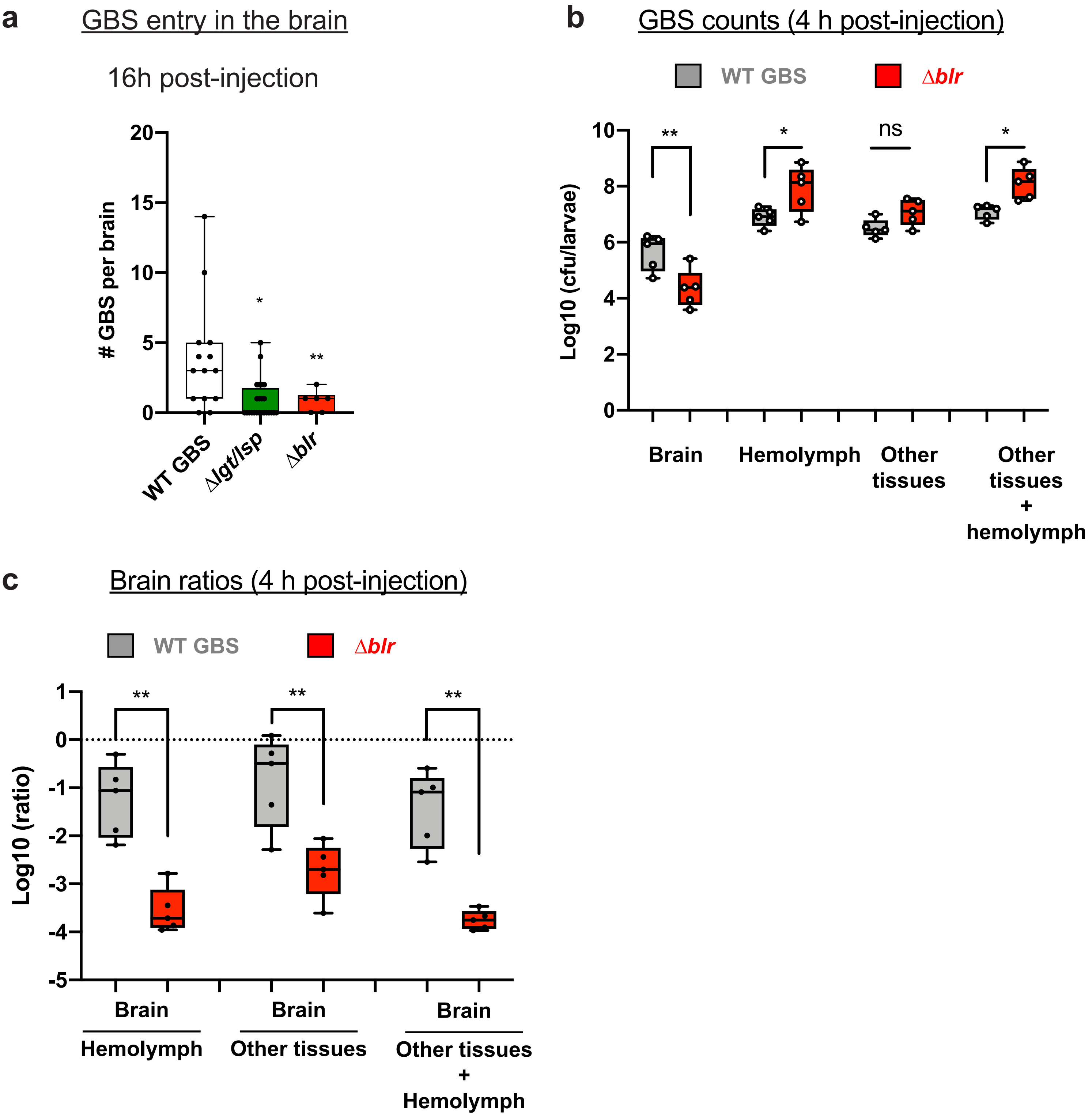

### Supplementary figure 6

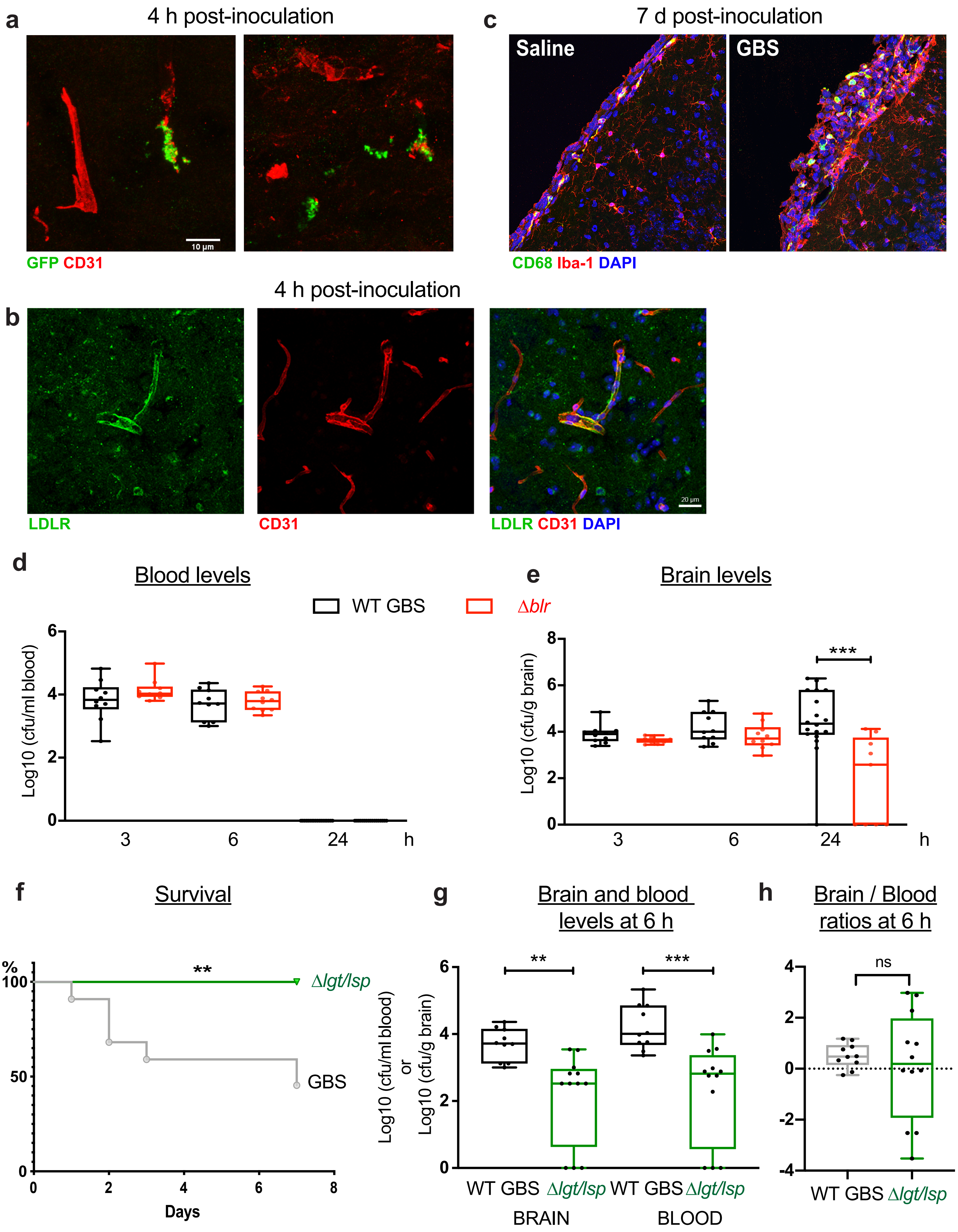
